# Non-monotonic recruitment of ventromedial prefrontal cortex during remote memory recall

**DOI:** 10.1101/202689

**Authors:** Daniel N. Barry, Martin J. Chadwick, Eleanor A. Maguire

## Abstract

Systems-level consolidation refers to the time-dependent reorganisation of memory traces in the neocortex, a process in which the ventromedial prefrontal cortex (vmPFC) has been implicated. Capturing the precise temporal evolution of this crucial process in humans has long proved elusive. Here, we used multivariate methods and a longitudinal functional MRI design to detect, with high granularity, the extent to which autobiographical memories of different ages were represented in vmPFC and how this changed over time. We observed an unexpected time-course of vmPFC recruitment during retrieval, rising and falling around an initial peak of 8-12 months, before re-engaging for older two and five year old memories. This pattern was replicated in two independent sets of memories. Moreover, it was further replicated in a follow-up study eight months later with the same participants and memories, where the individual memory representations had undergone their hypothesised strengthening or weakening over time. We conclude that the temporal engagement of vmPFC in memory retrieval seems to be non-monotonic, revealing a complex relationship between systems-level consolidation and prefrontal cortex recruitment that is unaccounted for by current theories.

**Author Summary:** Our past experiences are captured in autobiographical memories which allow us to recollect events from our lives long after they originally occurred. A part of the brain’s frontal lobe, called the ventromedial prefrontal cortex or vmPFC, is known to be important for supporting autobiographical memories especially as memories become more remote. The precise temporal profile of the vmPFC’s involvement is unclear, yet this information is vital if we are to understand how memories change over time and the mechanisms involved. In this study we sought to establish the time-course of vmPFC engagement in the recollection of autobiographical memories while participants recalled memories of different ages during functional magnetic resonance imaging (fMRI). Using a method that detects brain activity patterns associated with individual memories, we found that memory-specific neural patterns in vmPFC became more distinct over the first few months after a memory was formed, but then this initial involvement of vmPFC subsided after one year. However, more remote memories (two years and older), appeared to re-engage vmPFC once again. This temporal profile is difficult to accommodate within any single existing theory. Consequently, our results provoke a re-think about how memories evolve over time and the role played by the vmPFC.

## Introduction

We possess a remarkable ability to retrieve, with ease, one single experience from a lifetime of memories. How these individual autobiographical memories are represented in the brain over time is a central question of memory neuroscience which remains unanswered.

Consolidation takes place on two levels which differ on both a spatial and temporal scale. On a cellular level, the stabilisation of new memory traces through modification of synaptic connectivity takes only a few hours [1], and is heavily dependent upon the hippocampus [2–5]. On a much longer timescale, the neocortex integrates new memories, a form of consolidation termed “systems-level” [6]. The precise timeframe of this process is unknown. A related long-standing debate which has contributed to this uncertainty is whether or not the hippocampus ever relinquishes its role in autobiographical memory retrieval. One theory asserts that the hippocampus is not involved in the retrieval of memories after they have become fully consolidated to the neocortex [7]. Alternate views maintain that vivid, detailed autobiographical memories retain a permanent reliance on the hippocampus for their expression [8–12].

An undisputed feature of systems-level consolidation, however, is the strengthening of neural representations in the neocortex over time. Clarity on the time course of systems-level consolidation is therefore more likely to be achieved through scrutiny of its neocortical targets. While theoretical accounts often fail to specify these cortical locations, animal experiments have consistently implicated the medial prefrontal cortex. While this region has been associated with the formation [13, 14] and recall of recently acquired memories [15–17], in rodents it appears to be disproportionately involved in the retrieval of memories learned weeks previously [18–26]. The dependency on this region, which emerges over time, is facilitated by post-learning activation [27] and structural changes [28–30].

The evolutionary expansion of prefrontal cortex in humans makes it challenging to make direct anatomical comparisons with rodents, but the ventromedial prefrontal cortex (vmPFC) has been proposed as a homologous site of long-term memory consolidation [31]. It may appear surprising that an association between impaired autobiographical memory retrieval and vmPFC lesions has only recently started to be more precisely characterised [32]. However, there are a number of confounding factors in this field [33] - non-selectivity of vmPFC lesions, methodological differences in memory elicitation, and the tendency of patients with vmPFC damage to recollect events which have never occurred, a phenomenon known as confabulation [34].

Numerous functional MRI (fMRI) studies of vmPFC activity during autobiographical memory recall have been conducted, but with inconclusive results. Delay-dependent increases in retrieval-related activity have been observed in some studies [35, 36] but not others [37–39]. Autobiographical memory in particular induces robust vmPFC engagement [40] but it is unclear whether this activity increases [41], decreases [42], or remains constant in accordance with memory remoteness [43–52].

A powerful method of fMRI analysis which can help to bridge the empirical gap between the human and animal literatures is multi-voxel pattern analysis (MVPA), due to its increased sensitivity to specific neural representations [53]. Using this approach, Bonnici et al. [54] demonstrated that remote 10 year old autobiographical memories were more detectable in the vmPFC than recent two week old autobiographical memories, consistent with its proposed role as a long-term consolidation site. This difference was not apparent in other cortical areas, nor did it emerge from a standard univariate analysis. A follow up study two years later with the same participants and memories, demonstrated that the original two week old memories were now as detectable in the vmPFC as the remote memories [55]. This suggested the recent memories had been fully consolidated in the vmPFC after just two years, and perhaps even sooner.

The identification of this two year time window represented an opportunity to resolve the time course of systems-level consolidation with high precision. To do so, we sampled memories from four month intervals spanning a two year period, and compared their neural representations using fMRI. As opposed to the pattern classification approach employed by Bonnici et al. [54] to decode the neural signatures of individual memories, we used Representational Similarity Analysis (RSA) [56]. This method compares the consistency of neural patterns across repetitions of a single memory, against all other unrelated memories, to detect its unique informational content in a region of interest. Differences in the strength of memory representations across time periods were interpreted as delay-dependent engagement of the vmPFC. To verify observed time-sensitive differences, we followed the neural evolution of individual memories in a follow up study with the same participants and memories eight months later. The selection of numerous time-points characterised the consolidation process with unprecedented temporal resolution, while the longitudinal design was not only an opportunity to replicate these findings, but to observe systems-level consolidation in action.

Systems-level consolidation is generally assumed to be an incremental process, therefore, we considered a gradual linear trajectory of vmPFC recruitment as the most likely outcome. The alternative hypothesis was a rapid strengthening of vmPFC neural representations in the first few months after an event. The results conformed to neither scenario, and revealed an unexpected temporal relationship - a transient recruitment of the vmPFC beginning in the months following the initial experience, followed by an enduring signature of more remote memories. The second, longitudinal, experiment confirmed this finding. This is the first demonstration, to our knowledge, of such a temporal dissociation in vmPFC-mediated memory retrieval.

## Results

### Experiment 1

One week prior to the fMRI scan, with the assistance of personal photographs, participants (n=30) verbally recalled and rated the characteristics of autobiographical memories from eight time periods: memories that were 0.5 months old (0.5M, i.e., two week old memories), 4M, 8M, 12M, 16M, 20M, 24M and also 60M old - these latter memories serving as a definitive benchmark for remote (5 year old) memories (see Materials and methods, Fig 1A). Two memories from each time period which were sufficiently vivid, detailed, specific and unique in time and place were chosen for subsequent recall in the scanner. This meant that there were two full sets of memories. Participants created a short phrase pertaining to each autobiographical memory, which was paired with the photograph to facilitate recall during the subsequent fMRI scan.

**Fig 1.**
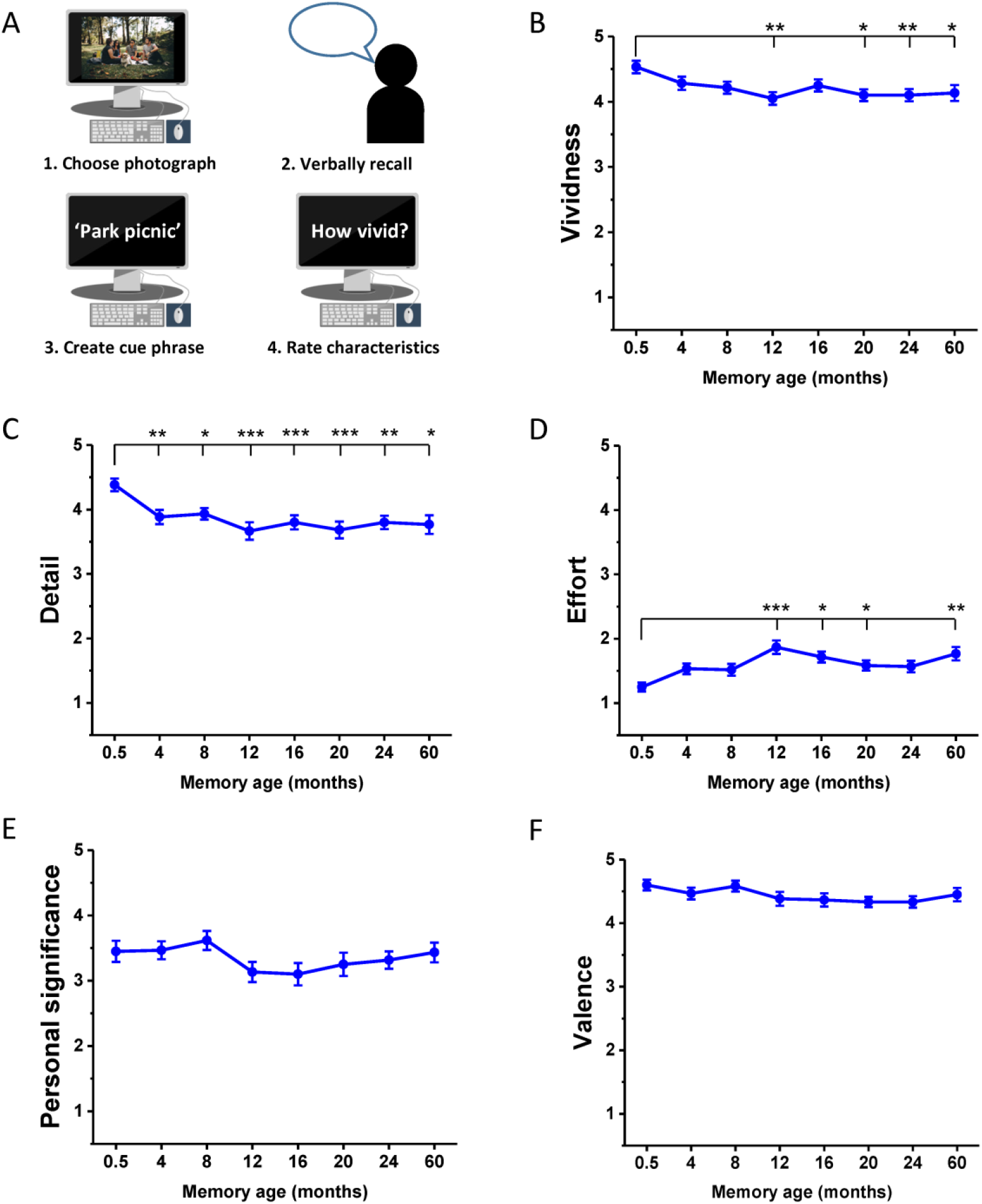
Memory harvesting and subjective ratings. (A) Schematic of the interview where the autobiographical memories were harvested. Participants recalled a memory which was cued by a personal photograph, chose a phrase to help remind them of this memory during the subsequent scanner task, and rated its characteristics. (B-F) Subjective ratings (means +/− 1SEM; see also means and SDs in Table A in S1 Table, and S1 Data for individual ratings across both sets of memories) of memory characteristics at each time period for Experiment 1, averaged across the two sets of memories. Ratings were on a scale of 1 to 5, where 1 was low and 5 was high. For emotional valence: 1-2 = negative, 3 = neutral, 4-5 = positive. * p < 0.05, ** p < 0.01, *** p < 0.001.

#### Comparable subjective recall phenomenology across memories

While all memories satisfied the criteria of being vivid and detailed, and the ratings were high (Fig 1; see means and SDs in Table A in S1 Table), subjective vividness nevertheless varied as a function of memory age (F_(7,203)_ = 3.45, p = 0.002), with the most recent, 0.5M old, memories rated higher than 12M (t_29_ = 4.08, p = 0.009), 20M (t_29_ = 3.88, p = 0.016), 24M (t_29_ = 4.18, p = 0.007) and 60M old memories (t_29_ = 3.45, p = 0.049, Fig 1B). Subjective ratings of detail also differed across time-points (F_(7,203)_ = 5.74, p < 0.001), once again the most recent 0.5M old memories were rated higher than 4M (t_29_ = 4.45, p = 0.003), 8M (t29 = 3.97, p = 0.012), 12M (t29 = 5.00, p < 0.001), 16M (t29 = 4.96, p < 0.001), 20M (t29 = 5.37, p < 0.001), 24M (t_29_ = 4.51, p = 0.003) and 60M old memories (t_29_ = 3.98, p = 0.012, Fig 1C). The expenditure of effort during recall also varied according to remoteness of memories (F_(7,203)_ = 5.79, p < 0.001), with 0.5M old memories being easier to recollect than 12M (t_29_ = −5.29, p < 0.001), 16M (t_29_ = −3.90, p = 0.015), 20M (t_29_ = −3.67, p = 0.027) and 60M old memories (t_29_ = −4.55, p = 0.003, Fig 1D). No significant difference was observed across time periods from 4M to 60M on any of these characteristics (all p > 0.05), nor did memories differ in their personal significance (F_(7,203)_ = 1.66, p = 0.120, Fig 1E) or emotional valence (F_(7,203)_ = 1.51, p = 0.166, Fig 1F) as a function of age.

In addition to these main ratings of interest, no difference was reported in the extent to which memories were recalled as active or static (F_(7,203)_ = 1.36, p = 0.224), or from a first or third person perspective (F_(3.69,107.02)_ = 1.09, p = 0.365) across time periods. The reported frequency at which memories were recalled since the original event (rated on a five point scale from “never” to “very frequently”), differed as a function of time (F_(5.11,148.04)_ = 4.36, p < 0.001), with the most recent 0.5M old memories thought about more frequently than 12M (t_29_ = 4.37 p = 0.004), 16M (t_29_ = 3.47, p = 0.046) and 24M (t_29_ = 3.71, p = 0.024) old memories (see S11 Data for individual ratings for these characteristics).

Overall, therefore, memories were generally well matched on subjective phenomenological ratings, satisfied the criteria of high quality of memory recall, with only small differences observed for the most recent 0.5M old memories compared to the other autobiographical memories, as might be expected.

#### Consistent level of details recalled across memories

To complement the subjective ratings of memory characteristics with a more objective assessment of their content, transcripts of participants’ memory interviews were scored using the Autobiographical Interview protocol ([57]; Materials and methods). In total for this first experiment, 10,187 details were scored. The mean (SD) number of internal details (bound to the specific ‘episodic’ spatiotemporal context of the event) and external details (arising from a general ‘semantic’ knowledge or references to unrelated events) are shown in Table B in S1 Table (see also Fig 2). They were then compared across time periods. In contrast to the subjective ratings of memory detail, the number of details recalled across memories from different time periods displayed only a non-significant trend (F_(4.54,131.66)_ = 1.92, p = 0.101). As expected, the number of internal and external details differed (F_(1,29)_ = 206.03, p < 0.001), with more internal details recalled for every time period (all p < 0.001). No interaction between time period and type of detail was observed (F_(7,203)_ = 1.87, p = 0.077). While a more targeted contrast of the most recent (0.5M) and most remote (60M) memories did reveal that 0.5M events contained more internal details (t(29) = 3.40, p = 0.002), this is consistent with participants’ subjective ratings, and implies that any observed strengthening of neural representations over time could not be attributable to greater detail at remote time-points. The number of external details recalled was remarkably consistent across all time periods, emphasising the episodic nature of recalled events irrespective of remoteness. Inter-rater reliabilities for the scoring (see Materials and methods) were high for both internal (ICC = 0.94) and external (ICC = 0.81) details.

**Fig 2.**
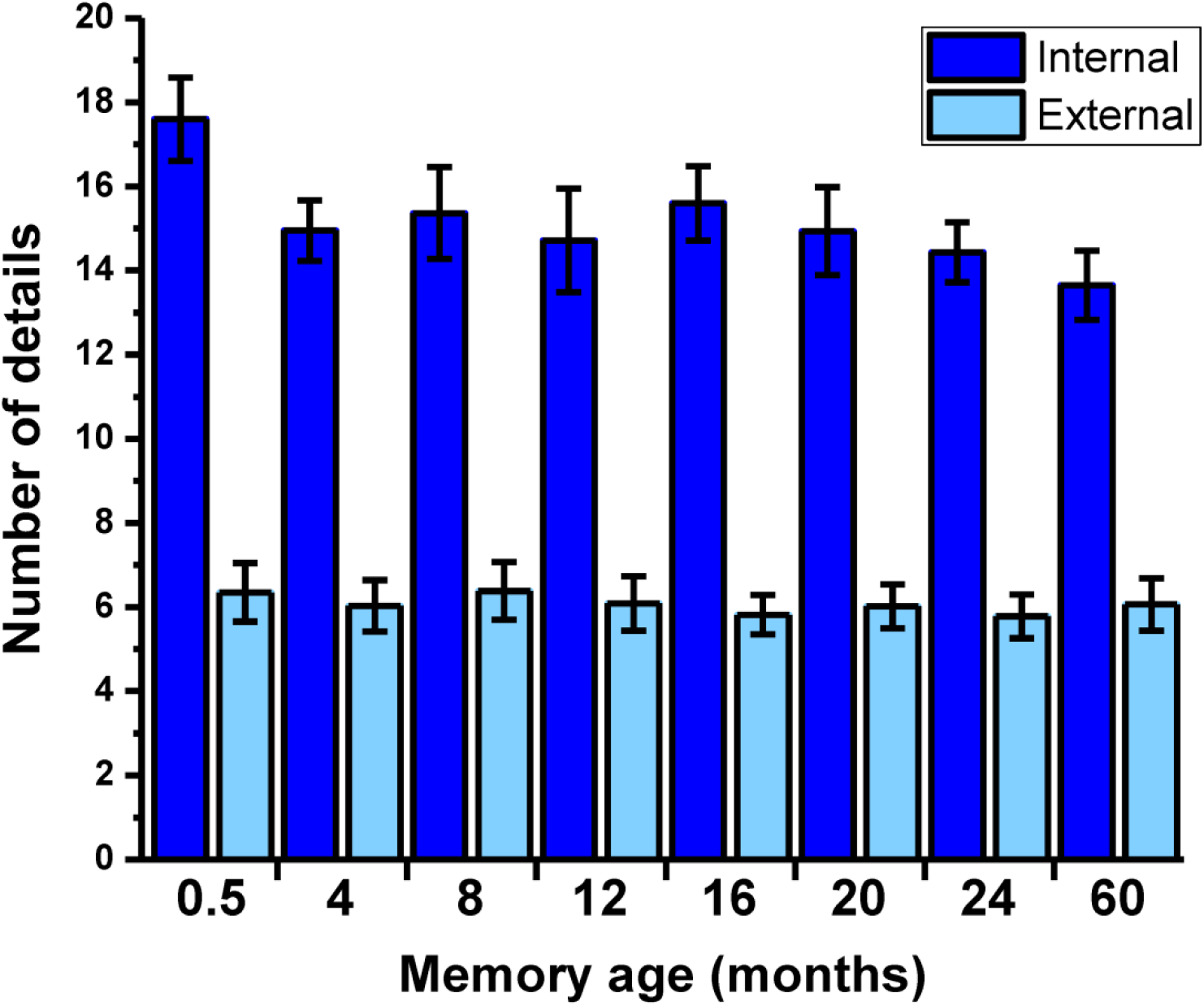
Objective scores for memory details. The mean +/− 1SEM (see also means and SDs in Table B in S1 Table, and S2 Data for individual participant scores) number of internal and external details at each time period, averaged across the two sets of autobiographical memories.

#### vmPFC engagement during recall was non-monotonic

vmPFC was delineated as the ventral medial surface of the frontal lobe and the medial portion of the orbital frontal cortex [58]. This comprises areas implicated in memory consolidation [31, 54, 55], namely Brodmann Areas 14, 25, ventral parts of 24 and 32, the caudal part of 10 and the medial part of BA 11 (Fig 3A, and Materials and methods).

**Fig 3.**
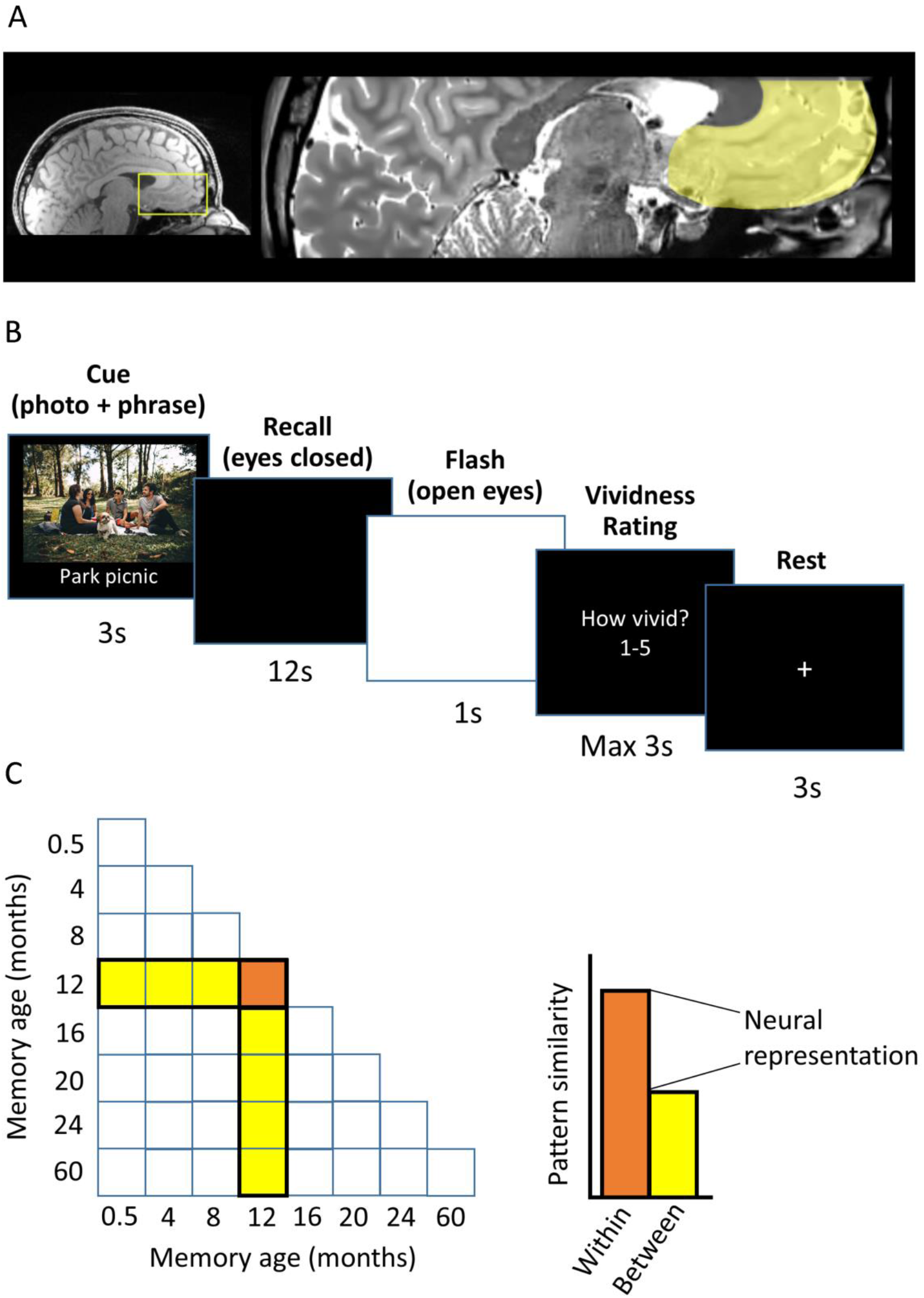
Experimental details. (A) The vmPFC is highlighted on an example participant’s structural MRI scan. (B) The timeline of an example trial from the scanning task. (C) Graphical illustration of the neural representation score calculation using RSA. The neural pattern similarity across trials recalling the same memory (orange) minus the mean pattern similarity between that memory and other memories (yellow) generates a “neural representation” score. A score significantly higher than zero indicates a neural pattern distinct to that memory is present in the vmPFC.

On each trial, the photograph and associated pre-selected cue phrase relating to each event were displayed on a screen for 3 seconds. Following removal of this cue, participants then closed their eyes and recalled the memory. After 12 seconds, the black screen flashed white twice, to cue the participant to open their eyes. The participant was then asked to rate how vivid the memory recall had been using a five-key button box, on a scale of 1-5, where 1 was not vivid at all, and 5 was highly vivid (Fig 3B).

We used RSA to quantify the extent to which the strength of memory representations in the vmPFC differed as a function of memory age. This was achieved by contrasting the similarity of neural patterns when recalling the same memory with their similarity to other memories to yield a “neural representation” score for each memory (see Materials and methods, Fig 3C). As there were two memories recalled per time period, the neural representation scores were averaged to produce one value for that time period.

We anticipated an increase in the strength of memory representations at some point between 0.5M and 24M, in line with the results of Bonnici and Maguire [55]. This is what we observed, where the most recent 0.5M memories were undetectable (t_29_ = 0.72, p = 0.477) in vmPFC, in contrast to the distinct neural signatures observed for 4M (t_29_ = 2.85, p = 0.008), 8M (t_29_ = 3.09, p = 0.004) and 12M (t_29_ = 3.66, p < 0.001) old memories (Fig 4A). These changes in the strength of memory representations were significant across time periods (F_(7,203)_ = 2.22, p = 0.034), with an observed increase in vmPFC recruitment from 0.5M to 8M (t_29_ = 2.07, p = 0.048) and 12M (t_29_ = −2.20, p = 0.036).

**Fig 4.**
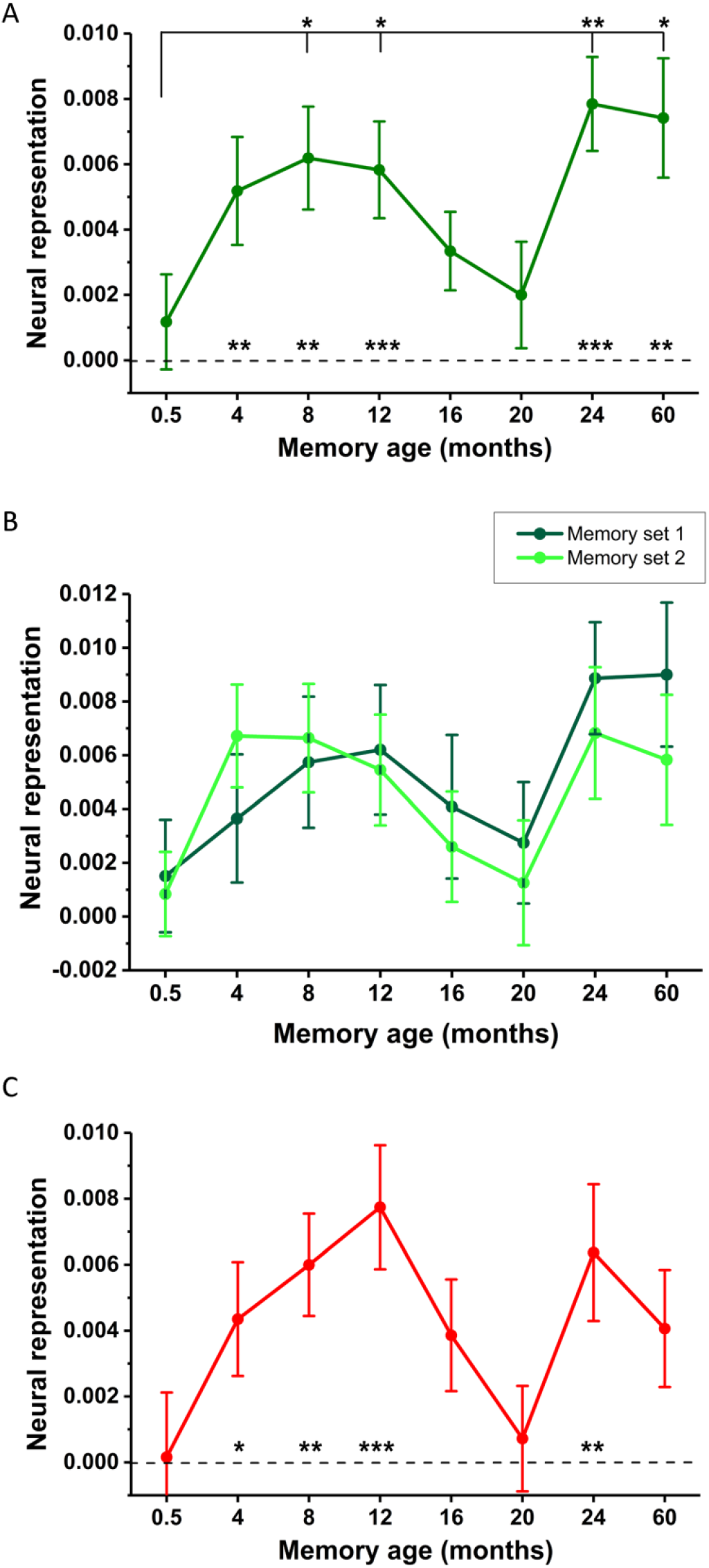
fMRI results of Experiment 1. (A) Mean +/− 1SEM neural representation scores at each time-point averaged across the two sets of memories. Asterisks above the dotted line indicate detectability of memories in vmPFC at each time-point. Asterisks above the solid line indicate significant increases in memory representations compared to the most recent (0.5M old) memories. * p < 0.05, ** p < 0.01, *** p < 0.001. See S1 Fig for the underlying representational similarity matrix and S2 Fig for a boxplot distribution of these data. (B) Neural representation scores at each time-point plotted separately for the two sets of autobiographical memories. (C) Neural representation scores when using a single identically-aged memory as a baseline. See S3 Data for individual participant scores.

However, what was observed for the following two time periods was unexpected - an apparent disengagement of the vmPFC over the next eight months as we observed weak detectability of memory representations in vmPFC for 16M (t_29_ = 1.85, p = 0.074) and 20M (t_29_ = 1.03, p = 0.310) old memories. Neither 16M (t_29_ = −1.06, p = 0.298) nor 20M memories (t_29_ = −0.40, p = 0.691) were more strongly represented than the recent 0.5M old memories. In contrast, the more remote 24M (t_29_ = 4.34, p < 0.001) and 60M (t_29_ = 3.55, p = 0.001) memories were detectable in the vmPFC, and significantly more so than the most recent memories (24M vs 0.5M, t_29_ = −2.93, p = 0.007; 60M vs 0.5M, t_29_ = −2.54, p = 0.017) as well as the more temporally proximal 20M old memories (24M vs 20M, t_29_ = −2.50, p = 0.018; 60M vs 20M, t _29_ = −2.32, p = 0.028).

The experimental design afforded us the opportunity to verify this non-monotonic pattern. As we sampled two memories per time-point, this time-dependent pattern should be evident in both sets of memories. As shown in Fig 4B, the two sets of memories followed a similar time-course of changes in representation within vmPFC. This is a compelling replication, given that the two memories from each time-period were unrelated in content as a prerequisite for selection, recalled in separate sessions in the scanner and analysed independently from each other.

The availability of two memories at each time-point also permitted the use of an alternative approach to calculating neural representation scores. Instead of using the similarity to memories from other time-points as a baseline, we could also assess if memories were similar to their temporally matched counterpart in the other set. As can be seen in Fig 4C, the non-monotonic pattern is preserved even when just using one identically aged memory as a baseline. In other words, the distinguishable patterns are specific to each individual memory rather than attributable to general retrieval processes associated with any memory of the same age.

An alternative explanation for memory representation scores which decreased over time is that the neural patterns became increasingly similar to memories from other time-points, rather than less consistent across repetitions, perhaps again reflecting more general retrieval processes. However as evident in S3 Fig, between-memory scores remained stable across all time-points, and did not differ in their statistical significance (F_(5.24,152.02)_ = 1.72, p = 0.13). If anything, there was a slight trend for higher between-memory scores to accompany higher within-memory scores. Therefore, the detectability of neural representations appeared to be driven by consistent within-memory neural patterns.

#### The observed temporal relationship is unique to vmPFC

Our main focus was the vmPFC, given previous work highlighting specifically this region’s role in representing autobiographical memories over time [54, 55]. We also scanned within a partial volume (to attain high spatial resolution with a reasonable TR), so were constrained in what other brain areas were available for testing (see Materials and methods). Nevertheless, we examined the same brain areas as Bonnici et al. [54], Bonnici and Maguire [55], additionally including the precuneus, given its role in autobiographical memory retrieval [59], and in no case did we observe a significant change in memory detectability across time periods - entorhinal/perirhinal cortex (F_(7,203)_ = 1.55, p = 0.154), hippocampus (F_(7,203)_ = 0.98, p = 0.445), posterior parahippocampal cortex (F_(7,203)_ = 1.41 p = 0.202), retrosplenial cortex (F_(7,203)_ = 0.69, p = 0.682), temporal pole (F_(7,203)_ = 1.78, p = 0.093), lateral temporal cortex (F_(4.86,141.03)_ = 0.68, p = 0.636) or precuneus (F_(7,203)_ = 0.789, p = 0.562). Of note, memories which were undetectable in the vmPFC were still represented in other brain regions at these time points (see S2 Table for neural representation score means and SDs, and S13 Data for individual participant scores). For example, 20 month old memories which did not appear to recruit the vmPFC during retrieval were represented in the majority of other regions comprising the core autobiographical memory network (precuneus, lateral temporal cortex, parahippocampal cortex, and approaching significance in the retrosplenial cortex (t_29_ = 1.83, p = 0.08)).

Following scanning, participants completed three additional ratings. They were asked to indicate the extent to which the memories were changed by the 6 repetitions during scanning on a scale ranging from 1 (not at all) to 5 (completely). They reported that the memories were not changed very much by repetition (mean: 2.61, SD: 0.74). They were also asked how often during scanning they thought about the memory interview one week previous on a scale of 1 (not at all) to 5 (completely), with participants indicating they rarely thought about the interview (mean: 2.29, SD: 1.01). Finally, participants were asked the extent to which the recall of memories from each time period unfolded in a consistent manner over the course of the session. A difference was observed (F_(7,203)_ = 2.78, p = 0.009), with the most recent 0.5M old memories being rated as more consistently recalled than the most remote 60M memories (t_29_ = 3.97, p = 0.012).

In addition to the region of interest (ROI)-based approach, a searchlight analysis was also conducted in MNI group normalised space to localise areas within the vmPFC where memories displayed high detectability across participants (see Materials and methods). We discovered a significant bilateral cluster of 652 voxels (see Fig A in S4 Fig), and subsequently used RSA to quantify the strength of neural representations at each time-point within this area (see Fig B in S4 Fig). The results were highly similar to the whole-ROI analysis in native space, suggesting the main result may be driven by more spatially confined activity within the vmPFC. However a searchlight approach is sub-optimal to answer the current research question, as it requires an *a priori* model RSM against which to compare the neural patterns at each searchlight sphere, whereas the ROI approach makes no such assumptions.

We also conducted a standard mass-univariate analysis on the whole volume with memory remoteness as a parametric regressor, and no area displayed either a significant increase or decrease in activity in accordance with memory age, consistent with the findings of Bonnici et al. [54]. In a similar parametric analysis, we did not find evidence of the modulation of univariate activity by in-scanner vividness ratings as might be suggested by the findings of Sheldon and Levine [60], however, all memories chosen for the current study were highly vivid in nature.

One concern when studying covert cognitive processes such as autobiographical memory in the fMRI scanner is participant compliance, because performance is subjectively reported rather than objectively assessed. However if participants were complying with task demands, there should be an association between in-scanner subjective ratings and the detectability of neural representations. When non-vivid trials were additionally incorporated into the RSA analysis, the mean memory representation score in the vmPFC for all participants averaged across time-points decreased from 0.0049 (SD 0.005) to 0.0044 (SD 0.005). In fact, the deleterious effect of including these extra non-vivid trials was evident in 24 out of the 30 participants. Such a consistent relationship between participants’ subjective ratings of their own memory performance and the sensitivity of the RSA analysis to detect memory representations, strongly suggests participants were performing the task as instructed.

### Rationale and predictions for Experiment 2

The non-monotonic pattern we observed in the fMRI data did not manifest itself in the subjective or objective behavioural data. In fact, the only difference in those data was higher ratings for the most recent 0.5M old memories. However, these were paradoxically the most weakly represented memories in the vmPFC, meaning the neural patterns were not driven by memory quality. The objective scoring of the memories confirmed comparable levels of detail provided for all memories, without any significant drop in episodic detail or increase in the amount of semantic information provided as a function of time. Therefore, the amount or nature of the memory details were not contributing factors.

Nevertheless, to verify that the results genuinely represented the neural correlates of memory purely as a function of age, one would need to study the effects of the passage of time on the individual neural representations. Therefore we invited the participants to revisit eight months later to recall the same memories again both overtly and during scanning; 16 of the participants agreed to return. In order to generate specific predictions for the neural representations during Experiment 2, we took the actual data for the 16 subjects from Experiment 1 who returned eight months later (Fig 5 green line, where the non-monotonic pattern is still clearly evident), and shifted it forwards by two time-points to simulate the expected pattern eight months later (Fig 5 pink dotted line). Note that for the 28M and 32M time periods in Experiment 2 we assumed they would have the same level of detectability as 24M old memories given the absence of data relating to these time periods from Experiment 1. We further assumed the neural representations between 60M and 68M would be unchanged.

**Fig 5.**
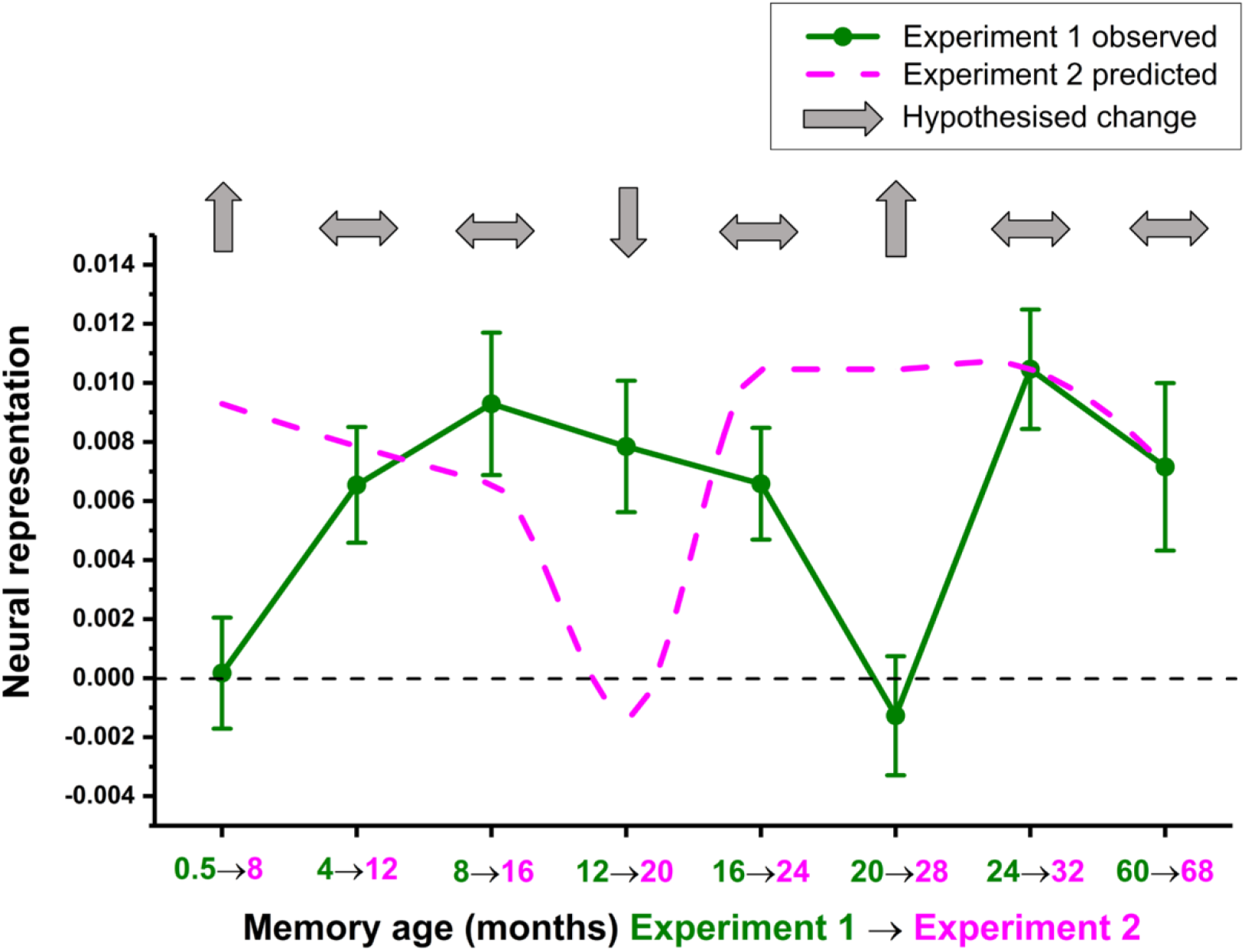
Predicted fMRI changes eight months later in Experiment 2. Predicted changes in the neural representations of individual autobiographical memories after eight months (pink dotted line), based on shifting the original observed data forward by two time-points for the 16 participants from Experiment 1 (green line) who returned for Experiment 2 (see S4 Data for original and predicted values). Light grey arrows indicate the hypotheses.

A comparison of the original and simulated neural representation scores yielded a number of clear hypotheses about how memory representations would change over time in the vmPFC. Two week old memories should become detectable eight months later, while the original 4M and 8M old memories should not differ in their representational strength. Twelve month old memories from Experiment 1 should be significantly less detectable, whereas 16M old memories should remain unchanged. The original 20M old memories should be better represented at 28M, whereas the 24 and 60 month old memories from Experiment 1 were not predicted to change over time.

### Experiment 2 (eight months later)

One week prior to the fMRI scan, with the assistance of the personal photographs and previously chosen phrases which were used as cues in Experiment 1, the participants verbally recalled and rated the characteristics of their autobiographical memories just as they had done eight months previously (see Materials and methods and Fig 6A).

**Fig 6.**
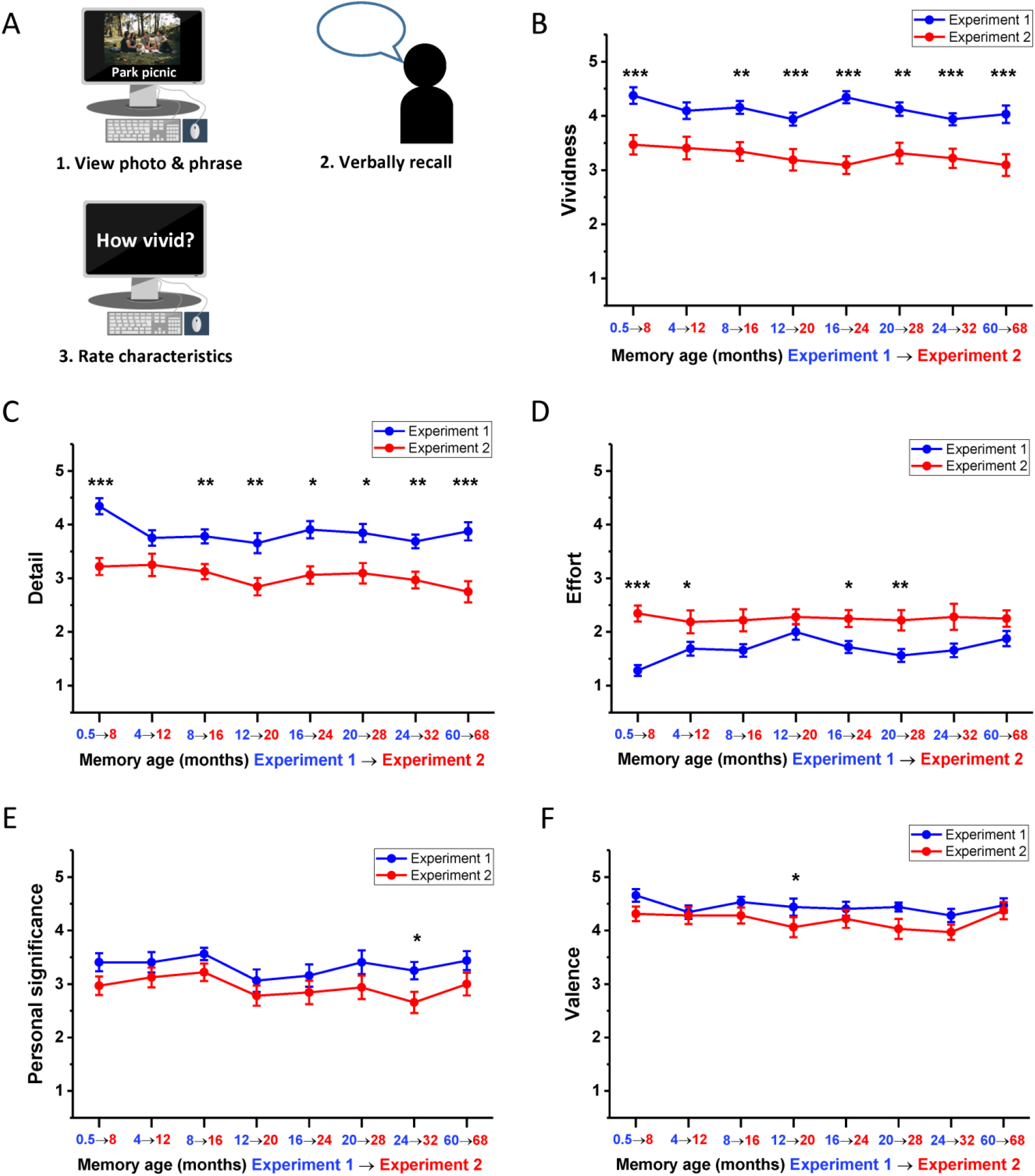
Memory recall and subjective ratings. (A) Schematic of the interview where participants recalled an autobiographical memory using their previously chosen photograph and cue phrase and rated its characteristics. (B-F) Subjective ratings (means +/− 1SEM; see also means and SDs in Table A in S1 Table and Table A in S3 Table) of memory characteristics at each time period for Experiment 1 (blue line, n=16 participants) and how the ratings of the same memories differed eight months later during Experiment 2 (red line, the same n=16 participants) averaged across the two sets of memories in both cases (see S5 Data for individual ratings across both sets of memories). Ratings were on a scale of 1 to 5, where 1 was low and 5 was high. For emotional valence: 1-2 = negative, 3 = neutral, 4-5 = positive. Asterisks indicate significant differences in memory ratings between Experiments 1 and 2; * p < 0.05, ** p < 0.01, *** p < 0.001.

#### Subjective ratings of phenomenology remain equivalent across memories

Means and SDs are provided in Table A in S3 Table. Autobiographical memories recalled during Experiment 2 did not differ across time periods on vividness (F_(7,105)_ = 0.83, p = 0.564), detail (F_(7,105)_ = 1.30, p = 0.257), effort (F_(7,105)_ = 0.11, p = 0.998), personal significance (F_(7,105)_ = 1.49, p = 0.180), valence (F _(7,105)_ = 1.06, p = 0.397), viewpoint (F_(3.42,51.22)_ = 1.24, p = 0.31) or motion (F_(3.95,59.32)_ = 1.43, p = 0.237). When asked how frequently they had thought about the autobiographical memories in the eight months between experiments (rated on a five point scale from “never” to “very frequently”), participants reported some change across time periods (F_(7,105)_ = 3.04, p = 0.006). However, the only significant difference between time periods was a lower recall frequency for now 32M old memories compared to the now 12M (t_15_ = 3.87, p = 0.042). Given the range of responses to this question across conditions (1.50-2.03), clearly participants had not given the memories much thought in the intervening eight months. Therefore, all memories recalled in Experiment 2 were extremely well matched in terms of their phenomenology, which reflects the consistency observed in ratings from eight months onwards in Experiment 1.

There were, however, differences in the absolute values of subjective ratings between the two experiments. There was a decrease in the reported vividness of all memories from Experiment 1 to Experiment 2 (F_(1,15)_ = 88.45, p < 0.001), from 0.5M to when they were 8M old (t_15_ = 6.21, p < 0.001), 8M to 16M (t_15_ = 4.21, p = 0.006), 12M to 20M (t_15_ = 5.48, p < 0.001), 16M to 24M (t_15_ = 7.07, p < 0.001), 20M to 28M (t_15_ = 4.10, p = 0.008), 24M to 32M (t_15_ = 5.97, p < 0.001) and 60M to 68M (t_15_ = 5.33, p < 0.001; Fig 6B). A comparable change was observed in the subjective impression of memory detail recalled following the eight month interlude (F_(1,15)_ = 126.81, p < 0.001), with a drop from 0.5M to 8M (t_15_ = 6.26, p < 0.001), 8M to 16M (t_15_ = 4.03, p = 0.009), 12M to 20M (t_15_ = 4.78, p = 0.002), 16M to 24M (t_15_ = 3.72, p = 0.016), 20M to 28M (t_15_ = 3.67, p = 0.018), 24M to 32M (t_15_ = 4.55, p < 0.003) and 60M to 68M (t_15_ = 9.67, p < 0.001; Fig 6C). Recalling memories eight months later was also perceived as more effortful (F_(1,15)_ = 43.32, p < 0.001), from 0.5M to 8M (t_15_ = −7.81, p < 0.001), 4M to 12M (t_15_ = −3.30, p = 0.039), 16M to 24M (t_15_ = −1.95, p = 0.021), and 20M to 28M (t_15_ = −4.03, p = 0.009; Fig 6D). The elapsed time between experiments also led to a reduction in the reported personal significance of memories (F_(1,15)_ = 11.82, p = 0.004), from 24M to 32M (t_15_ = 3.58, p = 0.022; Fig 6E). Ratings of emotional valence also changed over the eight month period (F_(1,15)_ = 9.78, p = 0.007), with a reported attenuation of the positivity of memories from 12M to 20M (t_15_ = 3.87, p = 0.012; Fig 6F). In addition to these main ratings, no difference was reported in the extent to which memories were recalled from a first or third person perspective (F_(1,15)_ = 0.513, p = 0.485) over the eight month period. The extent to which memories were recalled as active or static was altered by the passage of time between experiments (F_(1,15)_ = 11.01, p = 0.005), with the original 0.5M old memories becoming more static when 8M old (t15 = −3.42, p = 0.031). See S12 Data for individual ratings for these characteristics.

Despite the observed changes in some subjective ratings from Experiment 1 to Experiment 2, they were unidirectional across all time periods. As such, if the pattern of hypothesised emergence and disappearance of neural representations in vmPFC were to be supported in Experiment 2, then it could not be accounted for by changes in subjective ratings. Additionally, although the changes in subjective ratings across time tend to suggest a comparable degradation in memory quality across all time periods, this may be misleading. The ratings overall were still high, and these absolute changes in values could be influenced by participants’ expectations of their ability to recall memories after an extended period of time with high fidelity, because the objective scoring of memory detail revealed no such pattern, as we report in the next section.

#### A similar level of detail was recalled across experiments

As with Experiment 1, transcripts of participants’ memory interviews during Experiment 2 were scored using the Autobiographical Interview protocol ([57]; see Materials and methods)). A total of 6,444 details were scored (see Table B in S3 Table for means, SD). There was a difference in the number of details recalled across different time periods in Experiment 2 (F_(7,105)_ = 2.49, p = 0.021). However, this difference was only observed for external details (F_(7,105)_ = 3.25, p = 0.004), with more provided for 28M memories than 12M memories (t_15_ = −4.68, p = 0.008). As with Experiment 1, the number of internal and external details differed (F_(1,15)_ = 72.57, p < 0.001), with more internal details recalled for every time period (all p < 0.01). No interaction between time period and type of detail was observed (F_(7,105)_ = 0.87, p = 0.530).

When the objective scores for both experiments were compared, no significant difference was observed in the overall number of details provided eight months later (F_(1,15)_ = 1.93, p = 0.185; Fig 7). Furthermore, there was no significant interaction between experiment and time period (F_(1,15)_ = 1.97, p = 0.066), indicating that the amount of details provided for memories from any particular time period in Experiment 1 were not affected by the passage of time. Finally, no interaction was observed between experiment and type of detail provided (F_(1,15)_ = 2.27, p = 0.153), showing that the ratio of internal to external details was preserved across experiments.

**Fig 7.**
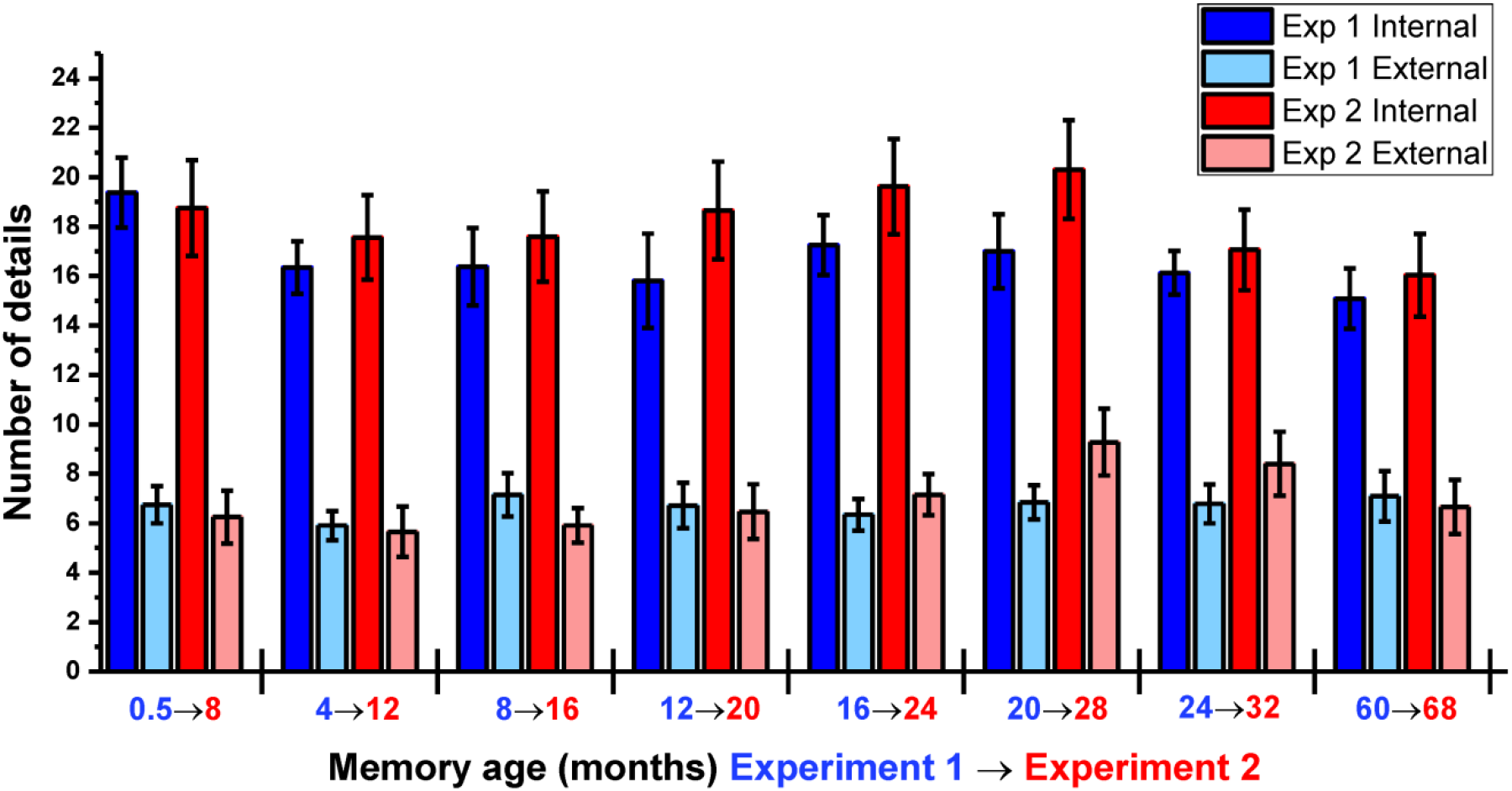
Objective scores for memory details over time. The mean +/− 1SEM (see also means and SDs in Table B in S1 Table and Table B in S3 Table) number of internal and external details at each time period for Experiment 1 (blue bars, n=16 participants) and Experiment 2 (red bars, the same n=16 participants), averaged across the two sets of autobiographical memories (see S6 Data for individual participant scores).

#### vmPFC memory representations undergo the predicted time-dependent changes

Participants were scanned in an identical fashion as Experiment 1 (see Materials and methods and Fig 3B), and neural representation scores for memories from each time point were again calculated.

When comparing the neural representation scores of memories from the eight original time periods in Experiment 1 with those of the same memories eight months later during Experiment 2, a main effect for experiment (F_(1,15)_ = 2.35, p = 0.146), or time period (F_(7,105)_ = 1.18, p = 0.323), was not observed, however, an interaction between experiment and time period emerged (F(7,105) = 3.46, p = 0.002). Closer examination via our planned comparisons (Fig 8A) revealed that seven out of the eight predictions made on the basis of the Experiment 1 findings were supported (Fig 8B). The original 0.5M old memories had increased in their representational strength in vmPFC eight months later (t_15_ = −1.84, p = 0.043), while the neural representation scores of the 4M and 8M old memories were essentially unchanged at 12M (t_15_ = 0.43, p = 0.677) and 16M (t_15_ = 1.22, p = 0.242) respectively. As expected, the original 12M old memories from Experiment 1 were eight months later more poorly represented in vmPFC when 20M months old (t_15_ = 1.85, p = 0.042). The original 16M old memories were unchanged in their representational strength at 24M (t_15_ = 1.38, p = 0.187), while 20M old memories were significantly more detectable in vmPFC at 28M (t_15_ = −2.69, p = 0.008). The most remote 60M memories did not differ in their neural representation scores eight months later (t_15_ = 0.86, p = 0.402). In fact the only finding which was inconsistent with the predictions generated by Experiment 1 was a decrease in the representation of 24M old memories when they were 32M of age (t_15_ = −2.69, p = 0.009). However, this prediction was based on the assumption that memories do not undergo further dynamic shifts in neural representation between two and five years, which may not be the case, and we did not have 32M data from Experiment 1 to corroborate this finding.

**Fig 8.**
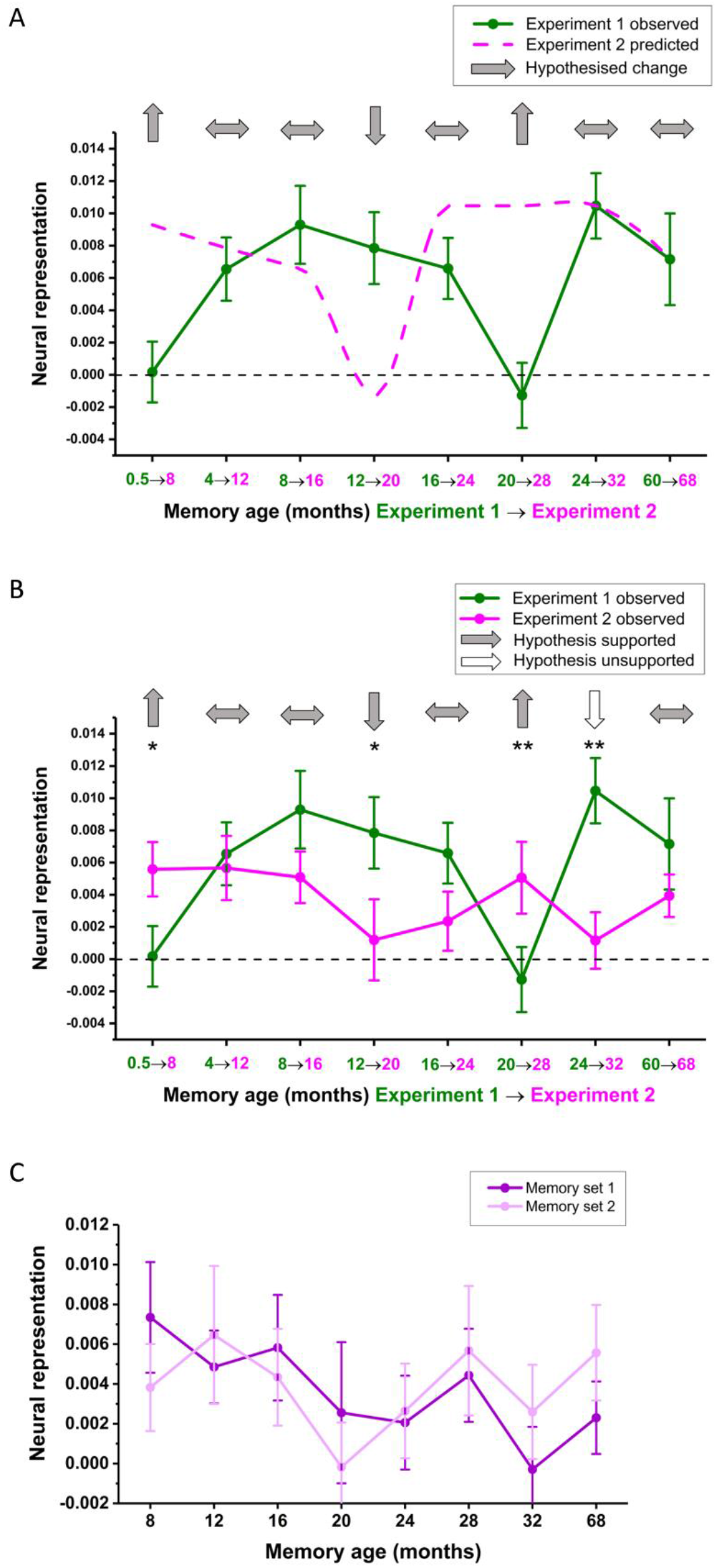
fMRI results of Experiment 2. (A) A reminder of the hypothesised changes in neural representations from Experiment 1 (green line) to Experiment 2 (pink line, reprinted from Fig 5). (B) Mean +/− 1SEM neural representation scores at each time-point averaged across the two sets of memories for Experiment 2 (pink line, n=16 participants) compared to the same memories from eight months previously (green line, the same n=16 participants). Light grey and white arrows indicate supported and unsupported hypotheses respectively; * p < 0.05, ** p < 0.01. (C) Neural representation scores at each time-point for Experiment 2, plotted separately for the two sets of autobiographical memories. See S7 data for individual participant scores.

For completeness, Fig 8C plots the neural representation scores for the two sets of memories in Experiment 2. As previously observed in Experiment 1, the two sets of memories displayed a similar time-course in terms of their neural representations, despite being recalled in separate scanning sessions, in a randomised order and analysed separately. As with Experiment 1, when examining other brain areas within the partial volume in Experiment 2, in no case did we find a significant difference in memory detectability across time periods.

Following scanning in Experiment 2, participants completed three additional ratings. They were asked to indicate the extent to which the memories were changed by the 6 repetitions during scanning on a scale ranging from 1 (not at all) to 5 (completely). As in Experiment 1, they reported that the memories were not changed very much by repetition (mean: 2.56, SD: 0.81). They were also asked how often they thought of the experience of recalling the memories in Experiment 1 while performing the scanning task in Experiment 2 on a scale of 1 (not at all) to 5 (during every memory). Participants indicated they rarely thought about Experiment 1 (mean: 1.75, SD: 0.93). Finally, the consistency of recall across time periods during the scanning session did not differ in Experiment 2 (F_(7,105)_ = 0.59, p = 0.761) or between the two experiments (F_(1,15)_ = 0.12, p = 0.733; see also Table A in S1 Table and Table A in S3 Table).

## Discussion

This study exploited the sensitivity of RSA to detect not only the extent to which memories of different ages were represented in the vmPFC, but how these representations changed over time. During Experiment 1, we observed detectability in vmPFC for memories at 4M to 12M of age, which was also evident at 24M and 60M. As expected, recent 0.5M old memories were poorly represented in vmPFC in comparison. Curiously, however, the same lack of detectability in vmPFC was observed for memories that were 16M to 20M old. This pattern persisted across separate sets of memories and was replicated in a follow-up study eight months later with the same participants and memories. Behavioural data failed to account for these time-dependent representational changes in either experiment and other regions failed to show a significant change in memory representations over time. These findings are difficult to accommodate within any single theoretical account of long-term memory consolidation [9, 12, 61–63], as neocortical recruitment is generally assumed to involve an ascending linear trajectory. Consolidation has been characterised as fluid and continuous [64], but the non-monotonic vmPFC engagement observed here suggests additional complexity in its temporal recruitment.

### Possible mechanisms underlying non-monotonic vmPFC recruitment

Over the course of consolidation in this study, the vmPFC twice alternated between disengagement and engagement, indicative of four separate stages. Below we consider, based on the latest theoretical developments and empirical research on systems-level consolidation and vmPFC functioning, the time-dependent processes which could underlie such a non-monotonic pattern.

#### Less than one month: quiescence during the early stages of systems-level consolidation

During retrieval of memories less than one month of age, there was a notable absence of vmPFC recruitment. This is consistent with previous human studies showing weaker representations of recent memories using pattern classification [54], and lower overall levels of fMRI activity [41]. Similarly, in animals, prefrontal activation is reduced during recent memory recall [18, 19], with lesions of this region generally preserving recent memory retrieval [21, 24]. Although prefrontal cells may be ‘tagged’ for subsequent consolidation around the time of encoding [14], they remain functionally immature and do not appear to significantly contribute to memory retrieval at this stage [26].

#### Four months to one year: disambiguation of competing consolidated representations

Over the subsequent three time periods in this study, vmPFC memory representations progressively strengthened. This echoes the time-dependent increases in prefrontal cortex activity observed in animal studies during memory retrieval [19, 65], and the disruption of remote memory following prefrontal inactivation [19, 21, 24]. While it has been demonstrated that the interim replay of recent experiences in the prefrontal cortex [66] coincides with lasting structural changes which facilitate subsequent recall [29], it is still unclear how these consolidated representations are utilised during retrieval. One prominent hypothesis is that the prefrontal cortex suppresses irrelevant representations [67], with corresponding evidence in animals that context-inappropriate memories are triggered following prefrontal inactivation [68]. Similarly in humans, vmPFC damage impairs the ability to suppress inappropriate memories through confabulation [34], and produces a tendency to confuse memories which have taken place in different contexts [69]. This is of potential relevance to the four to twelve month time period identified in the current study, as people remember vastly more memories from the past year than more remote life periods [70]. Therefore, the demands on memory disambiguation (the ability to correctly select from among similar memories during retrieval) are significantly increased across this timescale. For example, in attempting to recall the specific events from a party one attended months previously, multiple contemporaneous experiences which involve the same people, or have taken place in a similar context could interfere with recollection. While humans possess a large capacity for real-world stimuli in recent memory, an abundance of stored competing representations can be detrimental to memory performance during retrieval [71]. Therefore, vmPFC recruitment at these time-points may reflect a suppression of distractor representations which are inappropriate to the current retrieval intentions. Importantly, this would be a memory-specific process which would generate a consistent neural pattern every time a particular experience is recalled.

#### Sixteen to 20 months: time-induced decay negates the need for disambiguation

The progressive vmPFC disengagement observed over the following eight months suggests the suppression of interfering memories becomes less of a necessity over this period. Forgetting is a key attribute of an optimally functioning memory system [72], and the number of autobiographical events individuals can recall has been shown to decrease substantially between one and two years, before levelling off [73]. Therefore, the reduction in availability of potentially interfering memories from this time period may relieve the vmPFC from its role in disambiguating them from memories which have persisted through the consolidation process. For example, one may return from a vacation with many memories which contain multiple overlapping features, but this will inevitably be reduced to a few distinct experiences as time goes on.

#### Two years to five years: the emergence of schematic representations

If disambiguation ceases to be an issue for older memories, the robust re-engagement of vmPFC at more remote time periods suggests locally consolidated representations come to be used in a different way to assist in recall. From a theoretical perspective, systems-level consolidation is no longer viewed as the time-limited stabilisation of a static engram. Rather, the passage of time and repeated retrieval is thought to generate an additional representation which can complement the original detailed memory [10]. This emergent representation is schematic in nature, with an emphasis on general rather than specific details, and forms part of a flexible memory system which adapts to current demands. The network hub which supports schematic representations, suggested by both animal [17] and human neuroimaging [74] studies, is the medial prefrontal cortex. Further evidence is provided by patients with vmPFC lesions, who are resistant to false memory effects because schemas which would conflate actually studied and similar unseen words are not activated during retrieval [75]. Therefore, it is likely that the nature of memory representations in the vmPFC transform over the course of consolidation to become more schematic in nature. Accordingly, the re-engagement of vmPFC activity at more remote time-points in this study could point to the deployment of memory-congruent schema to assist in retrieval. For example, the vivid recollection of a memory from five years previously will likely require reorientation to an increasingly unfamiliar environment, an altered social network, and a different personal mindset. This may be facilitated by a rapid instantiation of relevant schematic representations in the vmPFC to bias retrieval in posterior brain regions, as proposed by Gilboa and Marlatte [76]. The non-monotonic recruitment observed here may, therefore, reflect not just the consolidation of neural representations, but their evolution over time and, most importantly, the way in which they are used to facilitate precise and holistic recollection. Importantly, vmPFC engagement during recall likely reflects not just task-related recruitment, but also communication with the hippocampus and other neocortical regions.

### Relevance to systems-level consolidation theories

The current findings have potential implications for the two dominant theoretical perspectives on systems-level consolidation. Standard Consolidation Theory [7] predicts that the passage of time promotes the strengthening of neural representations in the neocortex, but the duration of this process in humans is poorly specified. The current results suggest this process is accomplished over a relatively fast timescale on the order of months. The alternative perspective on consolidation, Multiple Trace Theory and Transformation Hypothesis [10], posits that over time, consolidation promotes the emergence of schematic, gist-like representations in the neocortex, which complement the original detailed memory. The re-engagement of the vmPFC at two years in this study may reflect the emergence of these generalised representations to facilitate specific recall at more remote time-points. Therefore, the consolidation of new memories in the neocortex may be reasonably rapid, whereas the transformation of these engrams may take place over a much longer timescale.

Using an autobiographical memory paradigm to study consolidation is preferable to laboratory-based episodic memory tests by virtue of its ecological validity, availability of temporally distant stimuli, clinical significance and context-dependent equivalence to animal tasks. However, studying autobiographical memory carries with it potential confounds which can affect interpretation of results. Below we consider why these factors cannot account for our observations.

### Consistency of recall and forgetting

Older memories may yield a higher RSA score if they are more consistently recalled. Here, however, participants actually rated 0.5M memories as more consistently recalled than 60 month old memories. Older memories were not impoverished in detail when compared to the detail available for recent memories. Moreover, an inspection of interview transcripts across experiments revealed participants rarely offered new details for previous memories when retested, countering the suggestion that increased detectability of old memories may arise from the insertion of new episodic or semantic details [77]. The consistency in recalled detail across experiments could be attributable to participants recalling in Experiment 2 what they had said during Experiment 1. However whether or not participants remembered by proxy is irrelevant, as they still recalled the specific details of the original event, removing forgetting as a potential explanation of changes in neural patterns over time.

### The influence of repetition

Retrieving a memory initiates reconsolidation, a transient state where memories are vulnerable to interference [78, 79]. Therefore, repeated retrieval may cause this process to have an influence on neural representations. However, all memories were recalled one week before the fMRI scan, so if such an effect was present it would be matched across time-points. Retrieval at this stage may also accelerate consolidation [80], yet if this was a major influence, we would likely have found 0.5M memories to be more detectable than they were. Further repetition of memories within the scanner in Experiment 1 took place over a timescale that could not affect consolidation processes or interpretation of the initial neural data. Nevertheless, this could arguably affect vmPFC engagement over a longer period of time [81] and thus perturb the natural course of consolidation, influencing the results of Experiment 2. However, given that seven out of the eight specifically hypothesised temporally sensitive changes in neural representations were supported, an altered or accelerated consolidation time-course appears highly unlikely. Again, recall recency was matched in Experiment 2 by the memory interview, and recall frequency between experiments was low.

Taking a more general and parsimonious perspective, the ratings demonstrate that, naturally, all memories are recalled on an occasional basis (Table A in S1 Table), therefore it seems highly unlikely that a mere six repetitions within a scanning session would significantly alter the time course of systems-level consolidation. It should also be noted that successful detection of neural patterns relied on the specific content of each memory, rather than being due to generic time-related retrieval processes (S4 Fig). One alternative to the current two-experiment longitudinal design to limit repetition across experiments would be to have a different group of participants with different memories for the second experiment. However the strength of the current approach was the ability to track the transformation in neural patterns of the same memories over time.

### The effect of selection

An alternative interpretation of the time-sensitive vmPFC engagement is a systematic bias in the content of selected memories. For example, annual events coinciding across all participants, such as a seasonal holiday. However, recruitment took place over a period of five months in an evenly spaced manner, ensuring that such events did fall into the same temporal windows across participants. The occurrence of personal events such as birthdays was also random across participants. The use of personal photographs as memory cues also limited the reliance on time of year as a method for strategically retrieving memories. Furthermore, the nature of memory sampling was that unique, rather than generic, events were eligible, reducing the likelihood of events which were repeated annually being included. Memory detectability was high at 12 month intervals such as one, two and five years in this study, suggesting perhaps it is easier to recall events which have taken place at a similar time of year to the present. However this should have been reflected in behavioural ratings, and equivalently strong neural representations for recent memories, but neither was observed. Most importantly, if content rather than time-related consolidation was the main influence on memory detectability, then we would not have observed any change in neural representation scores from Experiment 1 to Experiment 2, rather than the hypothesised shifts which emerged.

A related concern is that memories across time differ in nature because they differ in availability. Successful memory search is biased towards recency, meaning there are more events to choose from in the last few weeks, than remote time periods. Here, this confound is circumvented by design, given that search was equivalently constrained and facilitated at each time-point by the frequency at which participants took photographs, which was not assumed to change in a major way over time. These enduring “snap-shots” of memory, located within tight temporal windows (see Materials and methods) meant that memory selection was not confounded by retrieval difficulty or availability. It could also be argued that selection of time-points for this study should have been biased towards recency given that most forgetting occurs in the weeks and months after learning. However, it is important to dissociate systems-level consolidation from forgetting, as they are separate processes which are assumed to follow different time-courses. Memory forgetting follows an exponential decay [82], whereas systems-level consolidation has generally been assumed, until now, to be gradual and linear [83]. Our study was concerned only with vivid, unique memories which were likely to persist through the systems-level consolidation process.

A further potential concern regarding memory selection is that recent and remote memories which are comprised of equivalent levels of detail must be qualitatively different in some way. For example, selected remote memories must have been highly salient at the time of encoding to retain such high levels of detail. However, the underlying assumption that individual memories invariably become detail impoverished over time does not necessarily hold. While the volume of memories one can recall decreases over time [84], the amount of details one can recall from individual consolidated memories can actually increase over a one year delay [85]. While generalised representations are thought to emerge over the course of consolidation, they do not necessarily replace the original detailed memories [10], and the equivalent level of detail provided by participants across the two experiments here would suggest that memory specificity can be preserved over time. Furthermore, the possibility that remote memory selection may still be biased towards more salient memories is rendered unlikely by the method of memory sampling employed here. Because memories were chosen only from available photographic cues, the salience of recent and remote events was determined at the time of taking the photograph, and not during experimentation. These photographs served as potent triggers of remote memories which were not necessarily more salient than recent memories, and which may not have otherwise come to mind using a free recall paradigm. In addition, one would expect more salient remote memories to score higher than recent memories on subjective ratings of vividness, personal significance or valence, but this was not the case. Therefore, stronger neural representations at more remote time-points were likely due to consolidation-related processes rather than qualitative difference between recent and remote experiences at the time of encoding.

### Value

Given that the medial prefrontal cortex is often associated with value and emotional processing [86], could these factors have influenced the current findings? Humans display a bias towards consolidating positive memories [87], and remembered information is more likely to be valued than that which is forgotten [88]. Activity in vmPFC during autobiographical memory recall has been found to be modulated by both the personal significance and emotional content of memories [89]. However, in the current two experiments, memories were matched across time periods on these variables, and the selection of memories through photographs taken on a day-to day basis also mitigated against this effect. In the eight months between experiments, memories either remained unchanged or decreased slightly in their subjective ratings of significance and positivity, suggesting that these factors are an unlikely driving force behind the observed remote memory representations in vmPFC. For example, if recent memories in Experiment 1 were not well-represented in vmPFC because they were relatively insignificant, there is no reason to expect them to be more so eight months later, yet their neural representation strengthened over time nonetheless.

### Relation to previous findings

A methodological discrepancy between this experiment and that conducted by Bonnici et al. [54], is the additional use of a photograph to assist in cueing memories. One possible interpretation of the neural representation scores is they represent a role for the vmPFC in the maintenance of visual working memory following cue offset. However, the prefrontal cortex is unlikely to contribute to maintenance of visual information [90]. Furthermore, if this was the driving effect behind neural representations here, the effect would be equivalent across time-periods, yet it was not.

There is, however, an obvious inconsistency between the findings of the current study and that of Bonnici, et al. [54]. Unlike that study, we did not detect representations of 0.5M old memories in vmPFC. It could be that the support vector machine classification-based MVPA used by Bonnici et al. [54] is more sensitive to detection of memory representations than RSA, however, the current study was not optimised for such an analysis because it necessitated an increased ratio of conditions to trials. Nonetheless, the increase in memory representation scores from recent to remote memories was replicated and additionally refined in the current study with superior temporal precision. One observation which was consistent with the Bonnici findings was the detection of remote memories in the hippocampus, which also supports theories positing a perpetual role for this region in the vivid retrieval of autobiographical memories [10, 12]. However, the weak detectability observed at more recent time points may reflect a limitation of the RSA approach employed here to detect sparsely encoded hippocampal patterns, which may be overcome by a more targeted subfield analysis [91].

There are, however, distinct advantages to the use of RSA over pattern classification MVPA. RSA is optimal for a condition-rich design as it allows for the relationships between many conditions to be observed. For example, in the current experiment, a visual inspection of the group RSA matrix (S1 Fig) does not reveal an obvious clustering of recent or remote memories which would indicate content-independent neural patterns related to general retrieval processes. The approach employed by Bonnici et al. [54] assessed the distinctiveness of memories within each time-point from each other in order to detect memory representations. Should the neural patterns of a single memory become more consistent over time, yet also more similar to memories of the same age due to generic time-dependent mechanisms of retrieval, pattern classification would fail to detect a representation where one is present. In the current study, however, the two can be assessed separately, revealing memories at each time-point become distinct from both memories of all other ages (Fig 4A) and identically aged memories (Fig 4C). The machine learning approach employed by Bonnici et al. [54] to decode memory representations also requires the division of data into ‘training’ and ‘testing’ sets to classify unseen neural patterns [53]. This reduces the number of trials available for analysis, which would have been suboptimal for the current design because it would have necessitated an increased number of conditions and fewer trials per memory, whereas this restriction is not a necessity for RSA. Finally, because the pattern classification approach used by Bonnici et al. [54] compared memories from each time-point directly to each other, they could not be analysed independently. In the current RSA design, the two sets of memories could be analysed separately from each other to ascertain if the temporal patterns could be replicated in an independent set of data. As is evident in Fig 4B, the non-monotonic pattern of vmPFC recruitment was present in both sets of memories. The suitability of each MVPA method, therefore, depends on the study design and the research questions being posed.

In the light of our hypotheses, Experiment 2 generated one anomalous finding. Twenty-four month old memories from Experiment 1 were no longer well represented eight months later. Why memories around 32M of age are not as reliant on vmPFC is unclear, but unlike other time-periods, we cannot verify this finding in the current experiment, as we did not sample 32M memories during Experiment 1.

### Summary

The current results revealed that the recruitment of the vmPFC during the expression of autobiographical memories depends on the exact stage of systems-level consolidation, and that retrieval involves multiple sequential time-sensitive processes. These temporal patterns were remarkably preserved across completely different sets of memories in one experiment, and closely replicated in a subsequent longitudinal experiment with the same participants and memories. These findings support the notion that the vmPFC becomes increasingly important over time for the retrieval of remote memories. Two particularly novel findings emerged. First, this process occurs relatively quickly, by four months following an experience. Second, vmPFC involvement after this time fluctuates in a highly consistent manner, depending on the precise age of the memory in question. Further work is clearly needed to explore the implications of these novel results. Overall, we conclude that our vmPFC findings may be explained by a dynamic interaction between the changing strength of a memory trace, the availability of temporally adjacent memories, and the concomitant differential strategies and schemas that are deployed to support the successful recollection of past experiences.

## Materials and methods

### Ethics statement

This study was approved by the local research ethics committee (University College London Research Ethics Committee, approval reference 6743/002). All investigations were conducted according to the principles expressed in the Declaration of Helsinki. Written informed consent was obtained for each participant.

### Experiment 1

#### Participants

Thirty healthy, right handed participants (23 female) took part (mean age 25.3, SD 3.5, range 21-32). All had normal or corrected-to-normal vision.

#### Memory interview and selection of autobiographical memories

Participants were instructed to select at least three photographs from each of eight time-points in their past (0.5M, 4M, 8M, 12M, 16M, 20M, 24M and 60M relative to the time of taking part in the experiment) which reminded them of vivid, unique and specific autobiographical events. The sampling was retrospective, in that the photographs were chosen from the participants’ pre-existing photograph collections and not prospectively taken with the study in mind. Highly personal, emotionally negative or repetitive events were deemed unsuitable. An additional requirement was that memories from the same time period should be dissimilar in content. For the four most recent time periods (0.5M-12M), the memories should have taken place within a temporal window two weeks either side of the specified date yielding a potential window of one month, for the next three time points (16M-24M), three weeks either side to allow a window of six weeks, and one month either side for the most remote time point (60M), giving a two month window. This graded approach was adopted to balance temporal precision with the availability of suitable memories at more remote time-points.

Participants were asked to describe in as much detail as possible the specific autobiographical memory elicited by a photograph. General probes were given by the interviewer where appropriate (e.g., “what else can you remember about this event?”). Participants were also asked to identify the most memorable part of the event which took place within a narrow temporal window and unfolded in an event-like way. They then created a short phrase pertaining to this episode, which was paired with the photograph to facilitate recall during the subsequent fMRI scan (Fig 1A). Participants were asked to rate each memory on a number of characteristics (see main text, Figs 1 and 6, S1 Table and S3 Table), and two memories from each time period which satisfied the criteria of high vividness and detail, and ease of recall were selected for recollection during the fMRI scan.

#### Behavioural analyses

The interview was recorded and transcribed to facilitate an objective analysis of the details, and the widely-used Autobiographical Interview method was employed for scoring [57]. Details provided for each memory were scored as either “internal” (specific events, temporal references, places, perceptual observations and thoughts or emotions) or “external” (unrelated events, semantic knowledge, repetition of details or other more general statements). To assess inter-rater reliability, a subset of sixteen memories (n=2 per time period) were randomly selected across 16 different subjects and scored by another experimenter blind to the aims and conditions of the study. Intra-class coefficient estimates were calculated using SPSS statistical package version 22 (SPSS Inc, Chicago, IL) based on a single measures, absolute-agreement, 2-way random-effects model.

As two memories per time period were selected for later recall in the scanner, behavioural ratings were averaged to produce one score per time period. Differences in subjective memory ratings across time periods were analysed using a one-way repeated measures ANOVA with Bonferroni-corrected paired t-tests. Differences in objective memory scores of internal and external details across time periods were analysed using a two-way repeated measures ANOVA with Bonferroni-corrected paired t-tests. A threshold of p < 0.05 was used throughout both experiments. All ANOVAs were subjected to Greenhouse-Geisser adjustment to the degrees of freedom if Mauchly’s sphericity test identified that sphericity had been violated.

#### Task during fMRI scanning

Participants returned approximately one week later (mean 6.9 days, SD 1) to recall the memories while undergoing an fMRI scan. Prior to the scan, participants were trained to recall each of the 16 memories within a 12 second recall period (as in Bonnici et al. [54], Bonnici and Maguire [55]), when cued by the photograph alongside its associated cue phrase. There were two training trials per memory, and participants were asked to vividly and consistently recall a particular period of the original event which unfolded across a temporal window matching the recall period.

During scanning, participants recalled each memory six times (6 trials x 16 memories = 96 trials). The two memories from each time period were never recalled together in the same session, nor was any one memory repeated within each session, resulting in 12 separate short sessions with eight trials in each, an approach recommended for optimal detection of condition-related activity patterns using MVPA [92]. Trials were presented in a random order within each session. On each trial, the photograph and associated pre-selected cue phrase relating to each event were displayed on screen for three seconds. Following removal of this cue, participants then closed their eyes and recalled the memory. After 12 seconds, the black screen flashed white twice, to cue the participant to open their eyes. The participant was then asked to rate how vivid the memory recall had been using a five-key button box, on a scale of 1-5, where 1 was not vivid at all, and 5 was highly vivid. When the least vivid trials were excluded, the mean number of trials (/6) selected for analysis from each time-point were as follows: 0.5M: 5.65 (SD 0.57), 4M: 5.50 (SD 0.56), 8M: 5.43 (SD 0.55), 12M: 5.50 (SD 0.63), 16M: 5.50 (SD 0.59), 20M: 5.43 (SD 0.65), 24M: 5.42 (SD 0.56), 60M: 5.23 (SD 0.69).

#### MRI data acquisition

Structural and functional data were acquired using a 3T MRI system (Magnetom TIM Trio, Siemens Healthcare, Erlangen, Germany). Both types of scan were performed within a partial volume which incorporated the entire extent of the ventromedial prefrontal cortex (Fig 3A).

Structural images were collected using a single-slab 3D T2-weighted turbo spin echo sequence with variable flip angles (SPACE) [93] in combination with parallel imaging, to simultaneously achieve a high image resolution of ^∼^500 μm, high sampling efficiency and short scan time while maintaining a sufficient signal-to-noise ratio (SNR). After excitation of a single axial slab the image was read out with the following parameters: resolution = 0.52 × 0.52 × 0.5 mm, matrix = 384 × 328, partitions = 104, partition thickness = 0.5 mm, partition oversampling = 15.4%, field of view = 200 × 171 mm 2, TE = 353 ms, TR = 3200 ms, GRAPPA x 2 in phase-encoding (PE) direction, bandwidth = 434 Hz/pixel, echo spacing = 4.98 ms, turbo factor in PE direction = 177, echo train duration = 881, averages = 1.9. For reduction of signal bias due to, for example, spatial variation in coil sensitivity profiles, the images were normalized using a prescan, and a weak intensity filter was applied as implemented by the scanner’s manufacturer. To improve the SNR of the anatomical image, three scans were acquired for each participant, coregistered and averaged. Additionally, a whole brain 3D FLASH structural scan was acquired with a resolution of 1 × 1 × 1 mm.

Functional data were acquired using a 3D echo planar imaging (EPI) sequence which has been demonstrated to yield improved BOLD sensitivity compared to 2D EPI acquisitions [94]. Image resolution was 1.5mm^3^ and the field-of-view was 192mm in-plane. Forty slices were acquired with 20% oversampling to avoid wrap-around artefacts due to imperfect slab excitation profile. The echo time (TE) was 37.30 ms and the volume repetition time (TR) was 3.65s. Parallel imaging with GRAPPA image reconstruction [95] acceleration factor 2 along the phase-encoding direction was used to minimize image distortions and yield optimal BOLD sensitivity. The dummy volumes necessary to reach steady state and the GRAPPA reconstruction kernel were acquired prior to the acquisition of the image data as described in Lutti et al. [94]. Correction of the distortions in the EPI images was implemented using B0-field maps obtained from double-echo FLASH acquisitions (matrix size 64×64; 64 slices; spatial resolution 3mm^3^; short TE=10 ms; long TE=12.46 ms; TR=1020 ms) and processed using the FieldMap toolbox available in SPM [96].

#### MRI data preprocessing

fMRI data were analysed using SPM12 (www.fil.ion.ucl.ac.uk/spm). All images were first bias corrected to compensate for image inhomogeneity associated with the 32 channel head coil [97]. Fieldmaps collected during the scan were used to generate voxel displacement maps. EPIs for each of the twelve sessions were then realigned to the first image and unwarped using the voxel displacement maps calculated above. The three high-resolution structural images were averaged to reduce noise, and co-registered to the whole brain structural scan. EPIs were also co-registered to the whole brain structural scan. Manual segmentation of the vmPFC was performed using ITK-SNAP on the group averaged structural scan normalised to MNI space. The normalised group mask was warped back into each participant’s native space using the inverse deformation field generated by individual participant structural scan segmentations. The overlapping voxels between this participant-specific vmPFC mask and the grey matter mask generated by the structural scan segmentation were used to create a native-space grey matter vmPFC mask for each individual participant.

#### Representational Similarity Analysis

Functional data were analysed at the single subject level without warping or smoothing. Each recall trial was modelled as a separate GLM, which comprised the 12 second period from the offset of the memory cue to just before the white flash which indicated to the participant they should open their eyes. Motion parameters were included as regressors of no interest. RSA [56], was performed using the RSA toolbox (http://www.mrc-cbu.cam.ac.uk/methods-and-resources/toolboxes/) and custom MATLAB (version R2014a) scripts. In order to account for the varying levels of noise across voxels which can affect the results of multivariate fMRI analyses, multivariate noise normalisation [98] was performed on the estimated pattern of neural activity separately for each trial. This approach normalises the estimated beta weight of each voxel using the residuals of the first-level GLM and the covariance structure of this noise. This results in the down-weighting of noisier voxels and a more accurate estimate of the task-related activity of each voxel.

The average number of voxels analysed in the vmPFC across the two sets of memories was 5252 (SD 1227). Whole ROI-based analysis was preferred to a searchlight approach which would involve comparing neural with model similarity matrices [99], as we did not have strong *a priori* hypothesis about changes in neural representations over time against which to test the neural data, nor did we want to make assumptions regarding the spatial distribution of informative voxels in the vmPFC.

As participants recalled two memories per time-point, the dataset was first split into two sets of eight time points, which were analysed separately using RSA. To characterise the strength of memory representations in the vmPFC, the similarity of neural patterns across recall trials of the same memory was first calculated using the Pearson product-moment correlation coefficient, resulting in a “within-memory” similarity score. Then the neural patterns of each memory were correlated with those of all other memories, yielding a “between-memory” similarity score. Both within- and between-memory correlations were performed on trials from separate runs. For each memory, the between-memory score was then subtracted from the within-memory score to provide a neural representation score (Fig 3C). This score was then averaged across the two memories at each time-point. Results for the left and the right hemispheres were highly similar, and therefore the data we report here are from the vmPFC bilaterally. A distinctive neural pattern associated with the recall of memories at each time period would yield a score significantly higher than zero, which was assessed using a one-sample t-test. Strengthening or weakening of memory representations as a function of remoteness would result in a significant difference in memory representation scores across time periods, and this was assessed using a one-way repeated measures ANOVA with post-hoc two-tailed paired t-tests. Error bars on graphs displaying neural representation scores were normalised to reflect within- rather than between-subject variability in absolute values, using the method recommended by Cousineau [100] for within-subjects designs. The range of values that we observed are entirely consistent with those in other studies employing a similar RSA approach in a variety of learning, memory and navigation tasks in a wide range of brain regions [101–110].

#### Searchlight analysis

An RSA searchlight analysis was conducted in normalised space, on multivariate noise-normalised data within the ROI. This approach selected every voxel within the ROI, and using a volumetric approach which is constrained by the shape of the ROI, expanded the area around that voxel until an area of 160 voxels was reached. Within each of these spheres, memories were correlated with themselves, and other memories, analogous to the standard ROI approach. Then the resulting neural RSM was correlated using Spearman’s rank correlation coefficient with a model RSM which consisted of ones along the diagonal and zeros on the off-diagonal. This model RSM was used to detect if individual memories were detectable across all time-points. For every voxel, the average correlation from every sphere it participated in was calculated, to generate a more representative score of its informational content. Parametric assumptions regarding the spatial distribution of unsmoothed data may not hold. Therefore we used statistical nonparametric mapping (SnPM13) on the resulting searchlight images. We used 10,000 random permutations, a voxel-level significance threshold of t=3, and a family-wise-error corrected cluster-wise threshold of p<0.05 within an ROI.

### Experiment 2

#### Participants

Sixteen of the 30 participants who took part in Experiment 1 returned to take part in Experiment 2 (14 female, mean age 24.7, SD 3.1, range 21-33) approximately eight months later (8.4 months, SD 1.2).

#### Memory interview

Participants were presented with the 16 photographs and cue phrases associated with the autobiographical memories in Experiment 1 and were asked to describe in as much detail as possible the specific event which they had recalled previously. General probes were given by the interviewer where appropriate (e.g. “what else can you remember about this event?”). The interviewer availed of summarised transcripts from Experiment 1 to verify the same memory and details were being recalled. Participants then rated each memory on the same characteristics assessed in Experiment 1. The memory interview during Experiment 2 was also recorded and transcribed.

#### Behavioural analyses

The analysis of subjective and objective ratings for Experiment 2 followed exactly the same procedure as Experiment 1. The extent to which subjective ratings for the same memory had changed between Experiment 1 and Experiment 2 was assessed using a two-way (experiment x time period) repeated measures ANOVA with Bonferroni-corrected paired t-tests. Differences in objective memory ratings across experiments were analysed using a two (experiment) x two (detail) x eight (time period) repeated measures ANOVA with Bonferroni-corrected paired t-tests.

#### Task during fMRI scanning

Participants returned approximately one week later for the fMRI scan (mean 5.5 days, SD 3.7). Prior to scanning, only one reminder training trial per memory was deemed necessary given the prior experience of performing the task in Experiment 1. The scanning task remained unchanged from Experiment 1, aside from the re-randomisation of trials within each session. When the least vivid trials were excluded, the mean number of trials (/6) selected for analysis from each time period were as follows: 8M: 5.94 (SD 0.25), 12M: 5.97 (SD 0.13), 16M: 5.88 (SD 0.29), 20M: 5.88 (SD 0.29), 24M: 5.94 (SD 0.25), 28M: 5.94 (SD 0.17), 32M: 5.84 (SD 0.40), 68M: 5.81 (SD 0.36).

#### MRI data acquisition

Structural and functional data were acquired using the same scanner and scanning sequences as Experiment 1. However the prior acquisition of the partial volume structural MRI scans negated the need to include these in the protocol of Experiment 2.

#### MRI data preprocessing

fMRI data were preprocessed using the same pipeline as Experiment 1, with the additional step of co-registering the functional scans of Experiment 2 to the structural scans of Experiment 1, which enabled the use of the vmPFC masks from Experiment 1. First-level GLMs of each recall trial were constructed in an identical manner to Experiment 1.

#### Representational Similarity Analysis

RSA of the Experiment 2 fMRI data was conducted in an identical manner to Experiment 1. The average number of voxels analysed in the vmPFC across the two sets of memories for all participants was 5228 (SD 1765). To ascertain whether the observed neural representation scores had changed between Experiments 1 and 2, a two-way (experiment x time period) repeated measures ANOVA was performed. To investigate if these changes mirrored the predictions generated by the original data, paired t-tests were performed between the neural representation scores for each memory from Experiment 1 and Experiment 2, one-tailed if there was a hypothesised increase or decrease.

## Acknowledgements

We thank David Bradbury and the Imaging Support team for technical assistance.

## Supporting Information

**S1 Table.** Behavioural data for Experiment 1 (mean, SD) - Experiment 1 (n=30).

| **A. Subjective ratings** | 0.5M | 4M | 8M | 12M | 16M | 20M | 24M | 60M |
| --- | --- | --- | --- | --- | --- | --- | --- | --- |
| Vividness | 4.53 (0.52) | 4.28 (0.55) | 4.22 (0.50) | 4.05 (0.53) | 4.25 (0.50) | 4.10 (0.50) | 4.10 (0.52) | 4.13 (0.67) |
| Detail | 4.38 (0.54) | 3.88 (0.61) | 3.93 (0.49) | 3.67 (0.75) | 3.80 (0.60) | 3.68 (0.71) | 3.80 (0.57) | 3.77 (0.78) |
| Effort | 1.25 (0.39) | 1.53 (0.45) | 1.52 (0.50) | 1.87 (0.57) | 1.72 (0.47) | 1.58 (0.44) | 1.57 (0.49) | 1.77 (0.57) |
| Personal significance | 3.45 (0.88) | 3.47 (0.75) | 3.62 (0.80) | 3.13 (0.85) | 3.10 (0.93) | 3.25 (0.98) | 3.32 (0.72) | 3.43 (0.83) |
| Valence | 4.60 (0.46) | 4.47 (0.51) | 4.58 (0.47) | 4.38 (0.60) | 4.37 (0.57) | 4.33 (0.44) | 4.33 (0.50) | 4.45 (0.58) |
| Active(1)/static event(2) | 1.20 (0.25) | 1.25 (0.31) | 1.22 (0.31) | 1.20 (0.28) | 1.18 (0.31) | 1.30 (0.39) | 1.25 (0.34) | 1.37 (0.37) |
| Self(1)/other perspective(2) | 1.03 (0.13) | 1.17 (0.36) | 1.13 (0.29) | 1.12 (0.25) | 1.08 (0.19) | 1.10 (0.24) | 1.10 (0.20) | 1.08 (0.27) |
| Recall frequency | 3.23 (0.63) | 2.78 (0.61) | 2.95 (0.66) | 2.55 (0.70) | 2.72 (0.61) | 2.88 (0.69) | 2.63 (0.66) | 2.83 (0.65) |
| Consistency | 4.28 (0.49) | 4.08 (0.76) | 3.83 (0.75) | 3.93 (0.54) | 4.02 (0.65) | 3.98 (0.53) | 3.85 (0.71) | 3.73 (0.69) |
| **B. Objective scores** |  |  |  |  |  |  |  |  |
| Internal details | 17.60 (5.42) | 14.95 (3.95) | 15.37 (5.96) | 14.72 (6.75) | 15.60 (4.84) | 14.93 (5.74) | 14.43 (3.89) | 13.65 (4.50) |
| External details | 6.35 (3.82) | 6.03 (3.34) | 6.38 (3.75) | 6.08 (3.53) | 5.82 (2.56) | 6.02 (2.83) | 5.78 (2.84) | 6.07 (3.42) |

**S2 Table.**
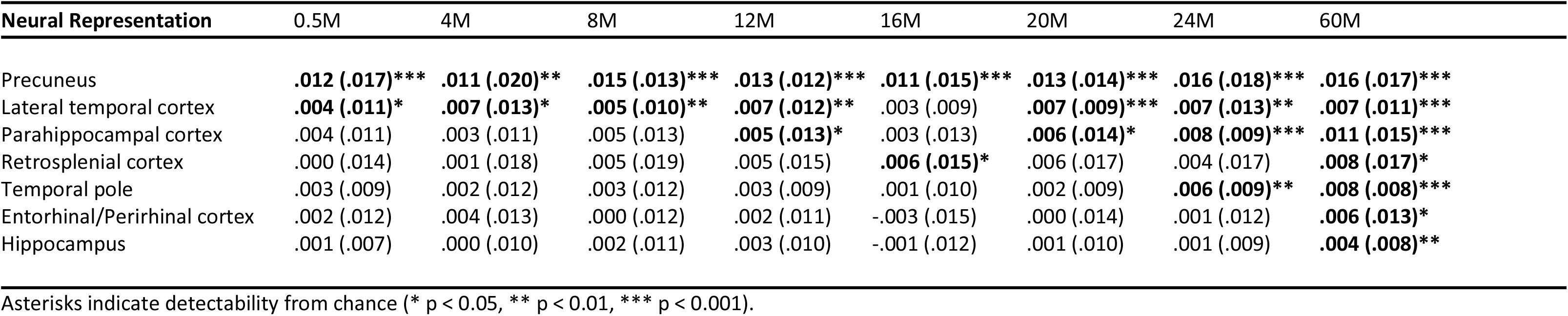
Neural representation scores (mean, SD) for other brain regions in Experiment 1 (n=30).

**S3 Table.**
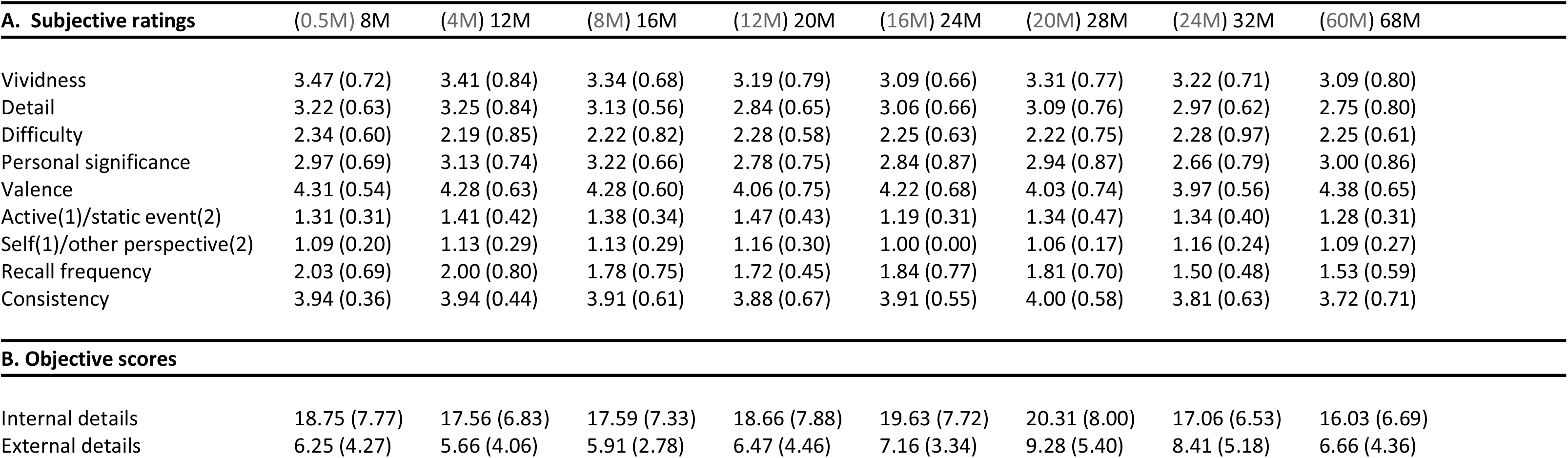
Behavioural data for Experiment 2 (mean, SD) - Experiment 2 (n=16).

| **A. Subjective ratings** | **(0.5M) 8M** | **(4M) 12M** | **(8M) 16M** | **(12M) 20M** | **(16M) 24M** | **(20M) 28M** | **(24M) 32M** | **(60M) 68M** |
| --- | --- | --- | --- | --- | --- | --- | --- | --- |
| Vividness | 3.47 (0.72) | 3.41 (0.84) | 3.34 (0.68) | 3.19 (0.79) | 3.09 (0.66) | 3.31 (0.77) | 3.22 (0.71) | 3.09 (0.80) |
| Detail | 3.22 (0.63) | 3.25 (0.84) | 3.13 (0.56) | 2.84 (0.65) | 3.06 (0.66) | 3.09 (0.76) | 2.97 (0.62) | 2.75 (0.80) |
| Difficulty | 2.34 (0.60) | 2.19 (0.85) | 2.22 (0.82) | 2.28 (0.58) | 2.25 (0.63) | 2.22 (0.75) | 2.28 (0.97) | 2.25 (0.61) |
| Personal significance | 2.97 (0.69) | 3.13 (0.74) | 3.22 (0.66) | 2.78 (0.75) | 2.84 (0.87) | 2.94 (0.87) | 2.66 (0.79) | 3.00 (0.86) |
| Valence | 4.31 (0.54) | 4.28 (0.63) | 4.28 (0.60) | 4.06 (0.75) | 4.22 (0.68) | 4.03 (0.74) | 3.97 (0.56) | 4.38 (0.65) |
| Active(1)/static event(2) | 1.31 (0.31) | 1.41 (0.42) | 1.38 (0.34) | 1.47 (0.43) | 1.19 (0.31) | 1.34 (0.47) | 1.34 (0.40) | 1.28 (0.31) |
| Self(1)/other perspective(2) | 1.09 (0.20) | 1.13 (0.29) | 1.13 (0.29) | 1.16 (0.30) | 1.00 (0.00) | 1.06 (0.17) | 1.16 (0.24) | 1.09 (0.27) |
| Recall frequency | 2.03 (0.69) | 2.00 (0.80) | 1.78 (0.75) | 1.72 (0.45) | 1.84 (0.77) | 1.81 (0.70) | 1.50 (0.48) | 1.53 (0.59) |
| Consistency | 3.94 (0.36) | 3.94 (0.44) | 3.91 (0.61) | 3.88 (0.67) | 3.91 (0.55) | 4.00 (0.58) | 3.81 (0.63) | 3.72 (0.71) |
| **B. Objective scores** |  |  |  |  |  |  |  |  |
| Internal details | 18.75 (7.77) | 17.56 (6.83) | 17.59 (7.33) | 18.66 (7.88) | 19.63 (7.72) | 20.31 (8.00) | 17.06 (6.53) | 16.03 (6.69) |
| External details | 6.25 (4.27) | 5.66 (4.06) | 5.91 (2.78) | 6.47 (4.46) | 7.16 (3.34) | 9.28 (5.40) | 8.41 (5.18) | 6.66 (4.36) |

**S1 Fig.**
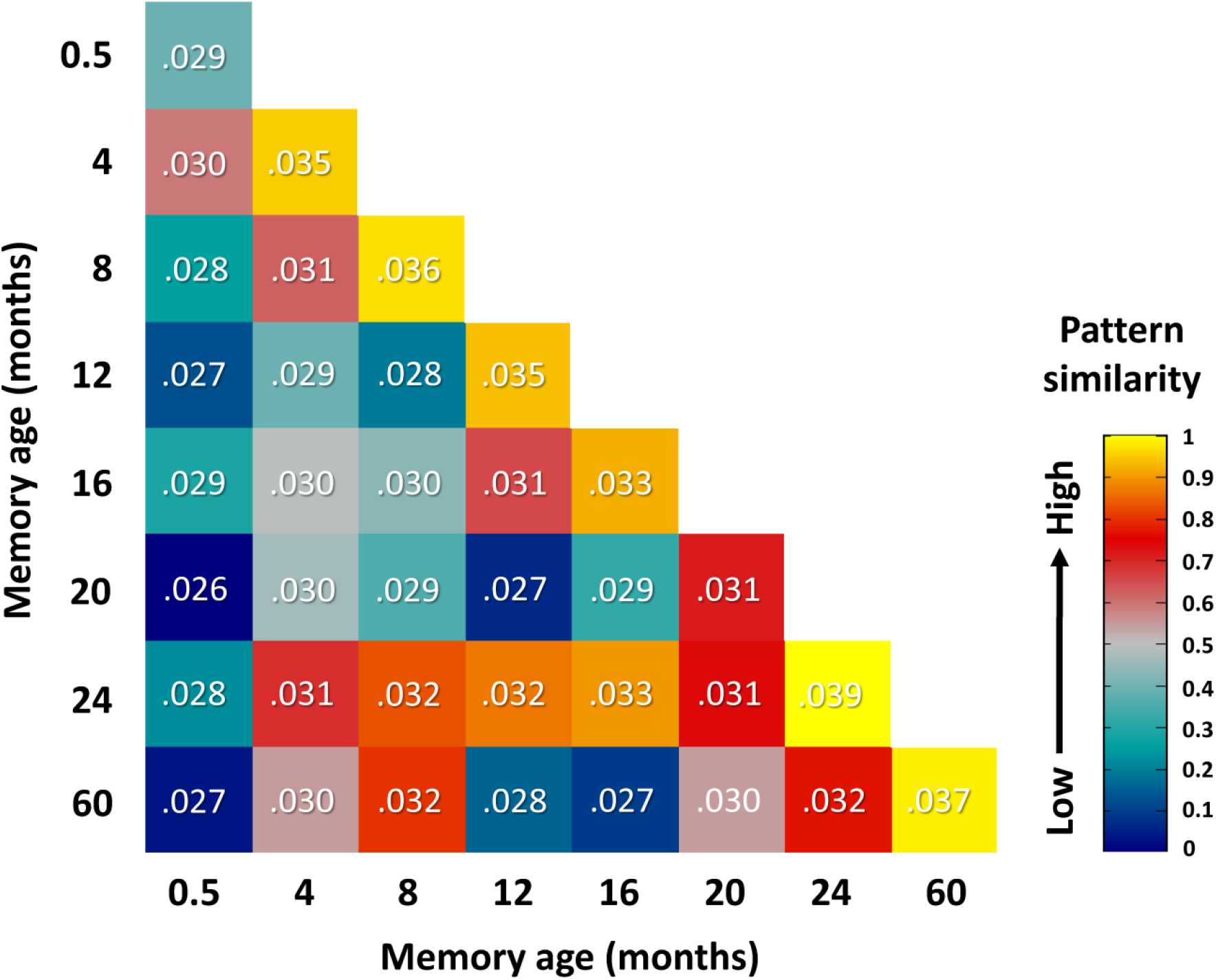
Representational similarity matrix of within- and between time-point pattern similarity values for Experiment 1. Each cell in this matrix contains the group mean pattern similarity score between memories from all sampled time-points, averaged across the two memory sets. The values along the diagonal represent the within-memory similarity for each time-point. Off-diagonal values indicate the correlation of neural patterns between memories of different ages, which are subsequently averaged to produce the baseline “between-memory” value and subtracted from the “within-memory” correlation to produce a neural representation score. For ease of visual inspection, all values are rank transformed, scaled between zero and one and colour coded to indicate the magnitude of pattern similarity.

**S2 Fig.**
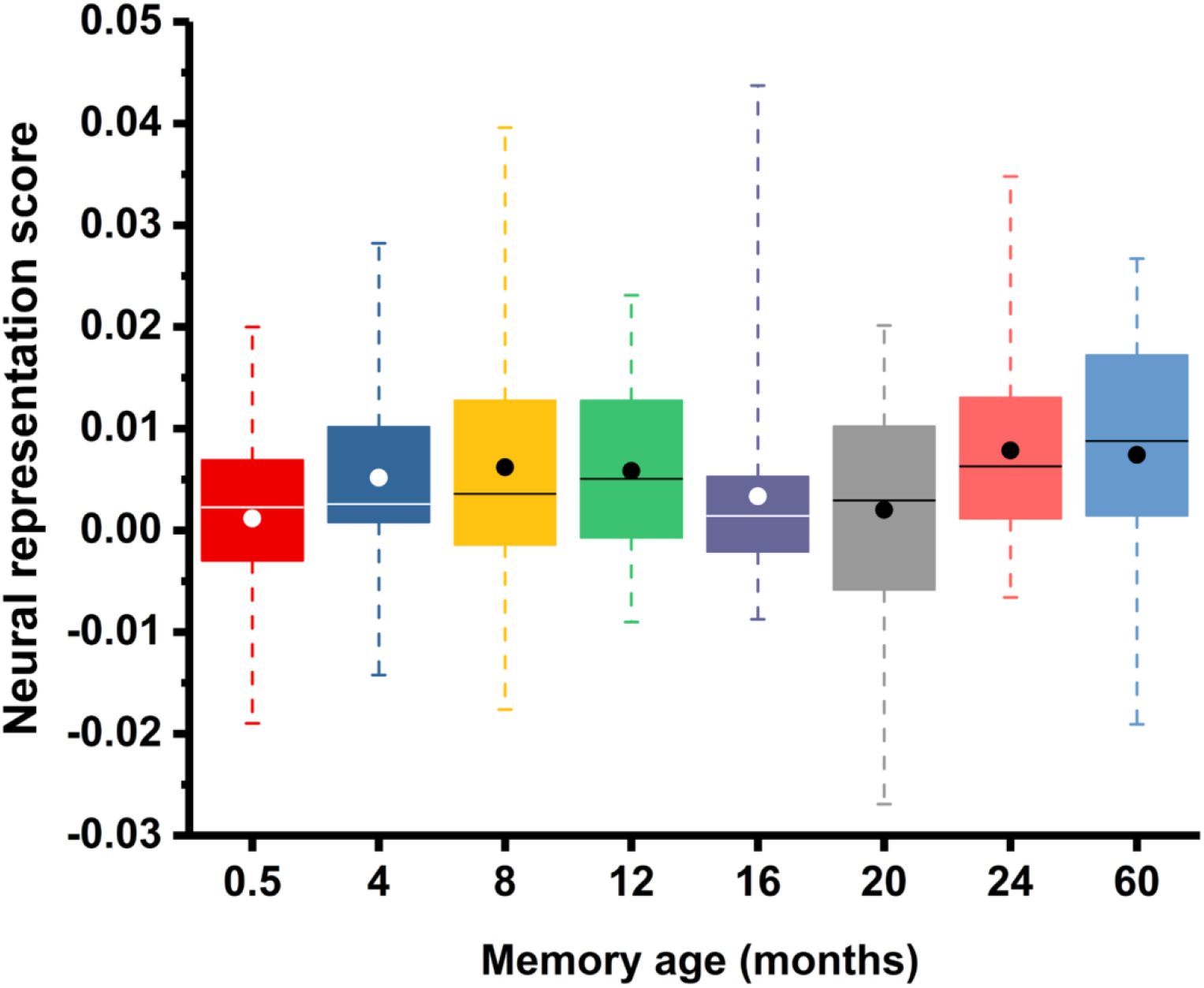
Boxplot of neural representation scores for Experiment 1. Boxes represent 25^th^ to 75^th^ percentiles around the median; whiskers represent minimum and maximum values, means are indicated by solid circles (see S8 Data for individual participant numerical values).

**S3 Fig.**
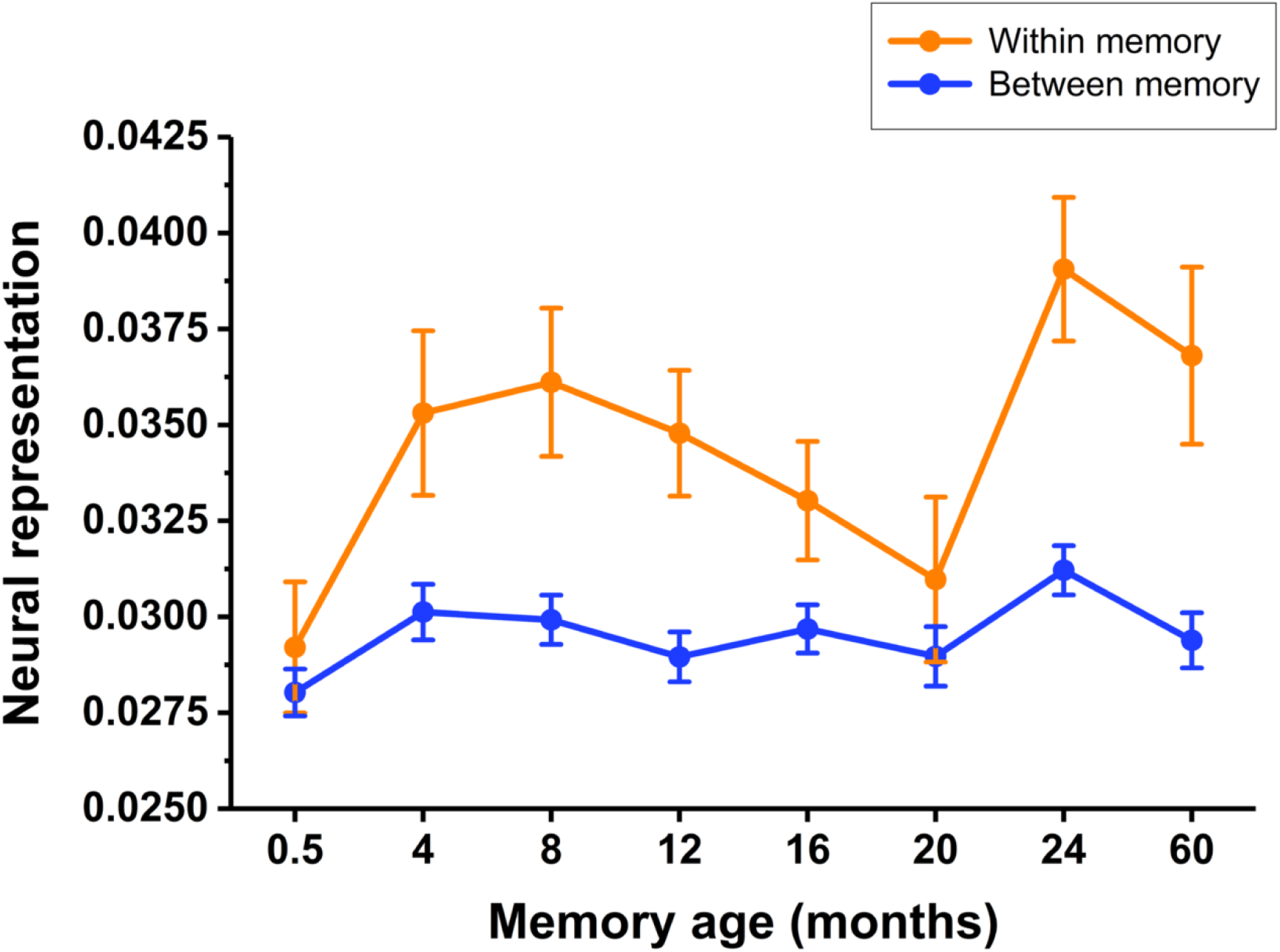
Within- versus between time-point pattern similarity for Experiment 1. Time-dependent changes in neural representation scores were driven by within- rather than between-memory scores (see S9 Data for individual participant numerical values).

**S4 Fig.**
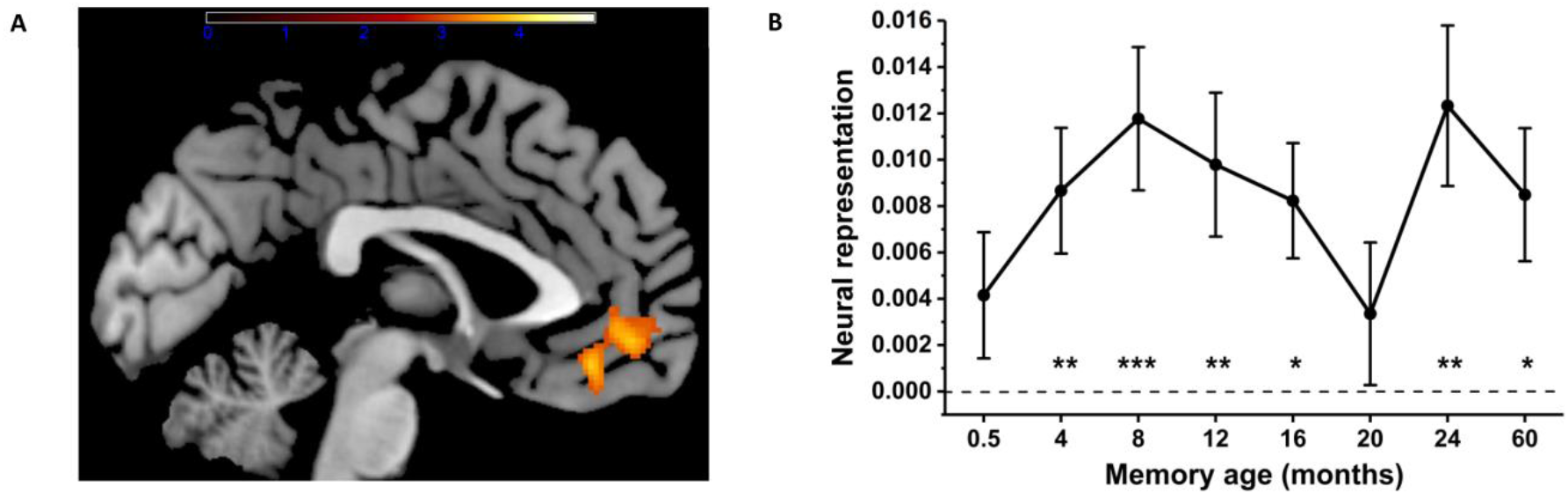
Results of the group vmPFC searchlight analysis in MNI space for Experiment 1. (A) Colour-coded areas represent the FWE-corrected T-statistic where within-memory detectability was higher than between-memory detectability across participants. (B) Comparison of memory detectability across time-points within this functionally defined area, showing highly similar results to the whole ROI analysis in native space (see S10 Data for individual participant numerical values).

## References

1. Kandel ER. The molecular biology of memory storage: a dialogue between genes and synapses. Science. 2001;294(5544):1030–8.

2. Pastalkova E, Serrano P, Pinkhasova D, Wallace E, Fenton AA, Sacktor TC. Storage of spatial information by the maintenance mechanism of LTP. Science. 2006;313(5790):1141–4.

3. Runyan JD, Dash PK. Inhibition of hippocampal protein synthesis following recall disrupts expression of episodic-like memory in trace conditioning. Hippocampus. 2005;15(3):333–9.

4. Guzowski JF. Insights into immediate-early gene function in hippocampal memory consolidation using antisense oligonucleotide and fluorescent imaging approaches. Hippocampus. 2002;12(1):86–104.

5. Morris RG, Moser EI, Riedel G, Martin SJ, Sandin J, Day M, et al. Elements of a neurobiological theory of the hippocampus: the role of activity-dependent synaptic plasticity in memory. Phil Trans R Soc Lond B, Biol Sci. 2003;358(1432):773–86.

6. Frankland PW, Bontempi B. The organization of recent and remote memories. Nature Rev Neurosci. 2005;6(2):119–30.

7. Squire LR, Genzel L, Wixted JT, Morris RG. Memory consolidation. Cold Spring Harbor Perspectives in Biology. 2015;7(8):a021766.

8. Moscovitch M, Rosenbaum RS, Gilboa A, Addis DR, Westmacott R, Grady C, et al. Functional neuroanatomy of remote episodic, semantic and spatial memory: a unified account based on multiple trace theory. J Anat. 2005;207(1):35–66.

9. Winocur G, Moscovitch M. Memory transformation and systems consolidation. JINS. 2011;17(5):766–80.

10. Moscovitch M, Cabeza R, Winocur G, Nadel L. Episodic memory and beyond: The hippocampus and neocortex in transformation. Ann Rev Psychol. 2016;67:105–34.

11. Maguire EA. Memory consolidation in humans: new evidence and opportunities. Exp Physiol. 2014;99(3):471–86.

12. Maguire EA, Mullally SL. The hippocampus: a manifesto for change. J Exp Psych: Gen. 2013;142(4):1180–9.

13. Zhao MG, Toyoda H, Lee YS, Wu LJ, Ko SW, Zhang XH, et al. Roles of NMDA NR2B subtype receptor in prefrontal long-term potentiation and contextual fear memory. Neuron. 2005;47(6):859–72.

14. Lesburgueres E, Gobbo OL, Alaux-Cantin S, Hambucken A, Trifilieff P, Bontempi B. Early tagging of cortical networks is required for the formation of enduring associative memory. Science. 2011;331(6019):924–8.

15. Leon WC, Bruno MA, Allard S, Nader K, Cuello AC. Engagement of the PFC in consolidation and recall of recent spatial memory. Learn Mem. 2010;17(6):297–305.

16. Einarsson EO, Nader K. Involvement of the anterior cingulate cortex in formation, consolidation, and reconsolidation of recent and remote contextual fear memory. Learn Mem. 2012;19(10):449–52.

17. Tse D, Takeuchi T, Kakeyama M, Kajii Y, Okuno H, Tohyama C, et al. Schema-dependent gene activation and memory encoding in neocortex. Science. 2011;333(6044):891–5.

18. Maviel T, Durkin TP, Menzaghi F, Bontempi B. Sites of neocortical reorganization critical for remote spatial memory. Science. 2004;305(5680):96–9.

19. Teixeira CM, Pomedli SR, Maei HR, Kee N, Frankland PW. Involvement of the anterior cingulate cortex in the expression of remote spatial memory. J Neurosci. 2006;26(29):7555–64.

20. Lopez J, Herbeaux K, Cosquer B, Engeln M, Muller C, Lazarus C, et al. Context-dependent modulation of hippocampal and cortical recruitment during remote spatial memory retrieval. Hippocampus. 2012;22(4):827–41.

21. Frankland PW, Bontempi B, Talton LE, Kaczmarek L, Silva AJ. The involvement of the anterior cingulate cortex in remote contextual fear memory. Science. 2004;304(5672):881–3.

22. Luo F, Zheng J, Sun X, Deng WK, Li BM, Liu F. Prelimbic cortex extracellular signal-regulated kinase 1/2 activation is required for memory retrieval of long-term inhibitory avoidance. Brain Res. 2017;1661:88–99.

23. Liu F, Zheng XL, Li BM. The anterior cingulate cortex is involved in retrieval of long-term/long-lasting but not short-term memory for step-through inhibitory avoidance in rats. Neurosci Lett. 2009;460(2):175–9.

24. Ding HK, Teixeira CM, Frankland PW. Inactivation of the anterior cingulate cortex blocks expression of remote, but not recent, conditioned taste aversion memory. Learn Mem. 2008;15(5):290–3.

25. Takehara K, Kawahara S, Kirino Y. Time-dependent reorganization of the brain components underlying memory retention in trace eyeblink conditioning. J Neurosci. 2003;23(30):9897–905.

26. Kitamura T, Ogawa SK, Roy DS, Okuyama T, Morrissey MD, Smith LM, et al. Engrams and circuits crucial for systems consolidation of a memory. Science. 2017;356(6333):73–8.

27. Takehara-Nishiuchi K, Nakao K, Kawahara S, Matsuki N, Kirino Y. Systems consolidation requires postlearning activation of NMDA receptors in the medial prefrontal cortex in trace eyeblink conditioning. J Neurosci. 2006;26(19):5049–58.

28. Restivo L, Vetere G, Bontempi B, Ammassari-Teule M. The formation of recent and remote memory is associated with time-dependent formation of dendritic spines in the hippocampus and anterior cingulate cortex. J Neurosci. 2009;29(25):8206–14.

29. Vetere G, Restivo L, Cole CJ, Ross PJ, Ammassari-Teule M, Josselyn SA, et al. Spine growth in the anterior cingulate cortex is necessary for the consolidation of contextual fear memory. Proc Natl Acad Sci USA. 2011;108(20):8456–60.

30. Bero AW, Meng J, Cho S, Shen AH, Canter RG, Ericsson M, et al. Early remodeling of the neocortex upon episodic memory encoding.Proc Natl Acad Sci USA. 2014;111(32):11852–7.

31. Nieuwenhuis IL, Takashima A. The role of the ventromedial prefrontal cortex in memory consolidation. Behavioural Brain Res. 2011;218(2):325–34.

32. Bertossi E, Tesini C, Cappelli A, Ciaramelli E. Ventromedial prefrontal damage causes a pervasive impairment of episodic memory and future thinking. Neuropsychologia. 2016;90:12–24.

33. McCormick C, Ciaramelli E, De Luca F, Maguire EA. Comparing and contrasting the cognitive effects of hippocampal and ventromedial prefrontal cortex damage: A review of human lesion studies. Neuroscience. 2017. doi: 10.1016/j.neuroscience.2017.07.066

34. Turner MS, Cipolotti L, Yousry TA, Shallice T. Confabulation: damage to a specific inferior medial prefrontal system. Cortex. 2008;44(6):637–48.

35. Takashima A, Petersson KM, Rutters F, Tendolkar I, Jensen O, Zwarts MJ, et al. Declarative memory consolidation in humans: a prospective functional magnetic resonance imaging study.Proc Natl Acad Sci USA. 2006;103(3):756–61.

36. Takashima A, Nieuwenhuis IL, Jensen O, Talamini LM, Rijpkema M, Fernandez G. Shift from hippocampal to neocortical centered retrieval network with consolidation. J Neurosci. 2009;29(32):10087–93.

37. Watanabe T, Kimura HM, Hirose S, Wada H, Imai Y, Machida T, et al. Functional dissociation between anterior and posterior temporal cortical regions during retrieval of remote memory. J Neurosci. 2012;32(28):9659–70.

38. Harand C, Bertran F, La Joie R, Landeau B, Mezenge F, Desgranges B, et al. The hippocampus remains activated over the long term for the retrieval of truly episodic memories. PloS One. 2012;7(8):e43495.

39. Furman O, Mendelsohn A, Dudai Y. The episodic engram transformed: Time reduces retrieval-related brain activity but correlates it with memory accuracy. Learn Mem. 2012;19(12):575–87.

40. Svoboda E, McKinnon MC, Levine B. The functional neuroanatomy of autobiographical memory: a meta-analysis. Neuropsychologia. 2006;44(12):2189–208.

41. Niki K, Luo J. An fMRI study on the time-limited role of the medial temporal lobe in long-term topographical autobiographic memory. J Cogn Neurosci. 2002;14(3):500–7.

42. Oddo S, Lux S, Weiss PH, Schwab A, Welzer H, Markowitsch HJ, et al. Specific role of medial prefrontal cortex in retrieving recent autobiographical memories: an fMRI study of young female subjects. Cortex. 2010;46(1):29–39.

43. Piolino P, Giffard-Quillon G, Desgranges B, Chetelat G, Baron JC, Eustache F. Re-experiencing old memories via hippocampus: a PET study of autobiographical memory. NeuroImage. 2004;22(3):1371–83.

44. Maguire EA, Henson RN, Mummery CJ, Frith CD. Activity in prefrontal cortex, not hippocampus, varies parametrically with the increasing remoteness of memories. Neuroreport. 2001;12(3):441–4.

45. Maguire EA, Frith CD. Lateral asymmetry in the hippocampal response to the remoteness of autobiographical memories. J Neurosci. 2003;23(12):5302–7.

46. Piefke M, Weiss PH, Markowitsch HJ, Fink GR. Gender differences in the functional neuroanatomy of emotional episodic autobiographical memory. Hum Brain Mapp. 2005;24(4):313–24.

47. Ryan L, Nadel L, Keil K, Putnam K, Schnyer D, Trouard T, et al. Hippocampal complex and retrieval of recent and very remote autobiographical memories: evidence from functional magnetic resonance imaging in neurologically intact people. Hippocampus. 2001;11(6):707–14.

48. Gilboa A, Winocur G, Grady CL, Hevenor SJ, Moscovitch M. Remembering our past: functional neuroanatomy of recollection of recent and very remote personal events. Cereb Cortex. 2004;14(11):1214–25.

49. Rekkas PV, Constable RT. Evidence that autobiographic memory retrieval does not become independent of the hippocampus: an fMRI study contrasting very recent with remote events. J Cogn Neurosci. 2005;17(12):1950–61.

50. Steinvorth S, Corkin S, Halgren E. Ecphory of autobiographical memories: an fMRI study of recent and remote memory retrieval. NeuroImage. 2006;30(1):285–98.

51. Soderlund H, Moscovitch M, Kumar N, Mandic M, Levine B. As time goes by: hippocampal connectivity changes with remoteness of autobiographical memory retrieval. Hippocampus. 2012;22(4):670–9.

52. Tsukiura T, Fujii T, Okuda J, Ohtake H, Kawashima R, Itoh M, et al. Time-dependent contribution of the hippocampal complex when remembering the past: a PET study. Neuroreport. 2002;13(17):2319–23.

53. Chadwick MJ, Bonnici HM, Maguire EA. Decoding information in the human hippocampus: a user's guide. Neuropsychologia. 2012;50(13):3107–21.

54. Bonnici HM, Chadwick MJ, Lutti A, Hassabis D, Weiskopf N, Maguire EA. Detecting representations of recent and remote autobiographical memories in vmPFC and hippocampus. J Neurosci. 2012;32(47):16982–91.

55. Bonnici HM, Maguire EA. Two years later - Revisiting autobiographical memory representations in vmPFC and hippocampus. Neuropsychologia. 2017 (Forthcoming: doi: 10.1016/j.neuropsychologia.2017.05.014.).

56. Kriegeskorte N, Kievit RA. Representational geometry: integrating cognition, computation, and the brain. Trends Cogn Sci. 2013;17(8):401–12.

57. Levine B, Svoboda E, Hay JF, Winocur G, Moscovitch M. Aging and autobiographical memory: Dissociating episodic from semantic retrieval. Psychol Aging. 2002;17(4):677–89.

58. Mackey S, Petrides M. Architecture and morphology of the human ventromedial prefrontal cortex. Eur J Neurosci. 2014;40(5):2777–96.

59. Hebscher M, Levine B, Gilboa, A. The precuneus and hippocampus contribute to individual differences in the unfolding of spatial representations during episodic autobiographical memory. Neuropsychologia. 2017;110:123–33.

60. Sheldon S, Levine B. Same as it ever was: vividness modulates the similarities and differences between the neural networks that support retrieving remote and recent autobiographical memories. NeuroImage. 2013;83:880–91.

61. Teyler TJ, DiScenna P. The role of hippocampus in memory: a hypothesis. Neurosci Biobehav Rev. 1985;9(3):377–89.

62. Squire LR, Alvarez P. Retrograde amnesia and memory consolidation: a neurobiological perspective. Curr Opin Neurobiol. 1995;5(2):169–77.

63. Nadel L, Winocur G, Ryan L, Moscovitch M. Systems consolidation and hippocampus: two views. Debates in Neuroscience. 2007;1(2-4):55–66.

64. Dudai Y. The restless engram: consolidations never end. Ann Rev Neurosci. 2012;35:227–47.

65. Barry DN, Coogan AN, Commins S. The time course of systems consolidation of spatial memory from recent to remote retention: A comparison of the Immediate Early Genes Zif268, c-Fos and Arc. Neurobiol Learn Mem. 2016;128:46–55.

66. Jadhav SP, Rothschild G, Roumis DK, Frank LM. Coordinated Excitation and Inhibition of Prefrontal Ensembles during Awake Hippocampal Sharp-Wave Ripple Events. Neuron. 2016;90(1):113–27.

67. Eichenbaum H. Memory: organization and control. Ann Rev Psychol. 2017;68:19–45.

68. Navawongse R, Eichenbaum H. Distinct pathways for rule-based retrieval and spatial mapping of memory representations in hippocampal neurons. J Neurosci. 2013;33(3):1002–13.

69. Schnider A, von Daniken C, Gutbrod K. Disorientation in amnesia. A confusion of memory traces. Brain. 1996;119 (Pt 5):1627–32.

70. Crovitz HF, Schiffman H. Frequency of episodic memories as a function of their age. Bull Psychonomic Soc. 1974;4:517–8.

71. Konkle T, Brady TF, Alvarez GA, Oliva A. Scene memory is more detailed than you think: the role of categories in visual long-term memory. Psychol Science. 2010;21(11):1551–6.

72. Hardt O, Nader K, Nadel L. Decay happens: the role of active forgetting in memory. Trends Cogn Sci. 2013;17(3):111–20.

73. Bauer PJ, Larkina M. Predicting remembering and forgetting of autobiographical memories in children and adults: a 4-year prospective study. Memory. 2016;24(10):1345–68.

74. vanKesteren MT, Beul SF, Takashima A, Henson RN, Ruiter DJ, Fernandez G. Differential roles for medial prefrontal and medial temporal cortices in schema-dependent encoding: from congruent to incongruent. Neuropsychologia. 2013;51(12):2352–9.

75. Warren DE, Jones SH, Duff MC, Tranel D. False recall is reduced by damage to the ventromedial prefrontal cortex: implications for understanding the neural correlates of schematic memory. J Neurosci. 2014;34(22):7677–82.

76. Gilboa A, Marlatte H. Neurobiology of schemas and schema-mediated memory. Trends Cogn Sci. 2017;21(8):618–31.

77. Berkers RM, van Kesteren MT. Autobiographical memory transformation across consolidation. J Neurosci. 2013;33(13):5435–6.

78. Nader K. Reconsolidation and the dynamic nature of memory. Cold Spring Harbor Perspectives in Biology. 2015;7(10):a021782.

79. Schwabe L, Wolf OT. New episodic learning interferes with the reconsolidation of autobiographical memories. PloS One. 2009;4(10):e7519.

80. Antony JW, Ferreira CS, Norman KA, Wimber M. Retrieval as a fast route to memory consolidation. Trends Cogn Sci. 2017;21(8):573–6.

81. Nadel L, Campbell J, Ryan L. Autobiographical memory retrieval and hippocampal activation as a function of repetition and the passage of time. Neural Plasticity. 2007;2007:90472.

82. Averell L, Heathcote A. The form of the forgetting curve and the fate of memories. J Math Psychol. 2011; 55(1):25–35.

83. McClelland JL, McNaughton BL, O'Reilly RC. Why there are complementary learning systems in the hippocampus and neocortex: Insights from the successes and failures of connectionist models of learning and memory. Psychol Rev. 1995;102(3):419–57.

84. Rubin DC. On the retention function for autobiographical memory. J Verb Learn Verb Be. 1982;21(1):21–38.

85. Campbell J, Nadel L, Duke D, Ryan L. Remembering all that and then some: recollection of autobiographical memories after a 1-year delay. Memory. 2011;19(4):406–15.

86. Grabenhorst F, Rolls ET. Value, pleasure and choice in the ventral prefrontal cortex. Trends Cogn Sci. 2011;15(2):56–67.

87. Anderson MC, Hanslmayr S. Neural mechanisms of motivated forgetting. Trends in Cogn Sci. 2014;18(6):279–92.

88. Rhodes MG, Witherby AE, Castel AD, Murayama K. Explaining the forgetting bias effect on value judgments: The influence of memory for a past test. Mem Cogn. 2017;45(3):362–74.

89. Lin WJ, Horner AJ, Burgess N. Ventromedial prefrontal cortex, adding value to autobiographical memories. Scientific Rep. 2016;6:28630.

90. Lee SH, Kravitz DJ, Baker CI. Goal-dependent dissociation of visual and prefrontal cortices during working memory. Nat Neurosci. 2013;16(8):997–9.

91. Bonnici HM, Chadwick MJ, Maguire EA. Representations of recent and remote autobiographical memories in hippocampal subfields. Hippocampus. 2013;23(10):849–54.

92. Coutanche MN, Thompson-Schill SL. The advantage of brief fMRI acquisition runs for multi-voxel pattern detection across runs. NeuroImage. 2012;61(4):1113–9.

93. Mugler JP, 3rd, Bao S, Mulkern RV, Guttmann CR, Robertson RL, Jolesz FA, et al. Optimized single-slab three-dimensional spin-echo MR imaging of the brain. Radiology. 2000;216(3):891–9.

94. Lutti A, Thomas DL, Hutton C, Weiskopf N. High-resolution functional MRI at 3 T: 3D/2D echo-planar imaging with optimized physiological noise correction. Magn Reson Med. 2013;69(6):1657–64.

95. Griswold MA, Jakob PM, Heidemann RM, Nittka M, Jellus V, Wang J, et al. Generalized autocalibrating partially parallel acquisitions (GRAPPA). Magn Reson Med. 2002;47(6):1202–10.

96. Hutton C, Bork A, Josephs O, Deichmann R, Ashburner J, Turner R. Image distortion correction in fMRI: A quantitative evaluation. NeuroImage. 2002;16(1):217–40.

97. Van Leemput K, Maes F, Vandermeulen D, Suetens P. Automated model-based tissue classification of MR images of the brain. IEEE Transactions on Medical Imaging. 1999;18(10):897–908.

98. Walther A, Nili H, Ejaz N, Alink A, Kriegeskorte N, Diedrichsen J. Reliability of dissimilarity measures for multi-voxel pattern analysis. NeuroImage. 2016;137:188–200.

99. Kriegeskorte N, Goebel R, Bandettini P. Information-based functional brain mapping. Proc Natl Acad Sci USA. 2006;103(10):3863–8.

100. Cousineau D. Confidence intervals in within-subject designs: A simpler solution to Loftus and Masson’s method. Tutorials in Quantitative Methods for Psychology. 2005;1(1):42–5.

101. Chadwick MJ, Jolly AEJ, Amos DP, Hassabis D, Spiers HJ. A goal direction signal in the human entorhinal/subicular region. Curr Biol. 2015;25(1):87–92.

102. Hsieh L-T, Gruber MJ, Jenkins LJ, Ranganath C. Hippocampal activity patterns carry information about objects in temporal context. Neuron. 2014;81(5):1165–78.

103. Schapiro AC, Turk-Browne NB, Norman KA, Botvinick MM. Statistical learning of temporal community structure in the hippocampus. Hippocampus. 2016;26(1):3–8.

104. Schuck Nicolas W, Cai Ming B, Wilson Robert C, Niv Y. Human orbitofrontal cortex represents a cognitive map of state space. Neuron. 2016;91(6):1402–12.

105. Kim M, Jeffery KJ, Maguire EA. Multivoxel pattern analysis reveals 3D place information in the human hippocampus. J Neurosci. 2017;37(16):4270–4279.

106. Hsieh L-T, Ranganath C. Cortical and subcortical contributions to sequence retrieval: Schematic coding of temporal context in the neocortical recollection network. NeuroImage. 2015;121:78–90.

107. Deuker L, Bellmund JLS, Navarro Schröder T, Doeller CF. An event map of memory space in the hippocampus. ELife. 2016;5:e16534.

108. Bellmund JLS, Deuker L, Navarro Schröder T, Doeller CF. Grid-cell representations in mental simulation. ELife. 2016;5:e17089.

109. Milivojevic B, Vicente-Grabovetsky A, Doeller CF. Insight reconfigures hippocampal-prefrontal memories. Curr Biol. 2015;25(7):821–30.

110. Staresina BP, Henson RNA, Kriegeskorte N, Alink A. Episodic reinstatement in the medial temporal lobe. J Neurosci. 2012;32(50):18150–6.

